# Vitamin B_6_ Catabolites Act as Interkingdom Signals Within the Marine Microbiome

**DOI:** 10.64898/2026.01.20.700305

**Authors:** Michael A. Ochsenkühn, Amin R. Mohamed, Lisa S.Y. Coe, Cong Fei, Sten Littman, Daniela Tienken, Justin R. Seymour, Jean-Baptiste Raina, Shady A. Amin

## Abstract

More than half of all net primary productivity in the oceans is exchanged and recycled among marine microbial communities, through a network of complex interactions that fuel the global carbon cycle and control oceanic biogeochemistry^1,2^. Cryptic chemical signals control these interactions, thereby governing interspecies metabolite exchanges and structuring the ocean microbiome^3^. Here, we show that vitamin B6 catabolites—molecules long assumed to be inert byproducts of metabolism—in fact serve as potent, bidirectional, and widely distributed chemical signals that structure metabolite exchanges between marine eukaryotic phytoplankton and bacteria. We demonstrate that a ubiquitous diatom and its bacterial symbiont exchange five intermediates and catabolites in the glycerophospholipid and vitamin B6 metabolic pathways, previously not implicated in interspecies interactions. Acting in combination, these signals induce broad transcriptional responses that prime each organism to anticipate metabolic input from its partner and enter a symbiotic-like state, despite having no measurable effect on growth. We further identify and biochemically validate the enzymes biosynthesizing two of these signals: diatom-derived 4-pyridoxolactone and bacterially-derived 4-pyridoxate—a catabolite long regarded as bacterial waste^4^. Genes encoding these enzymes are globally distributed and co-expressed across diverse sunlit ocean microbial communities, indicating that vitamin B6-mediated communication is widespread in the ocean. Together, these findings reveal an unexpected signaling role for vitamin B6 and demonstrate that canonical metabolic byproducts can function as information-rich signals structuring interkingdom interactions in the oceans.

---

Interactions between phytoplankton and bacteria underpin the global carbon cycle and shape marine biogeochemical cycles^5^. These interspecies interactions can span mutualism to parasitism^6^. During beneficial interactions, phytoplankton provide dissolved organic matter (DOM) to associated bacteria in exchange for vitamin cofactors, iron, hormones, and reduced nitrogen^5^. These reciprocal exchanges unfold within the phycosphere—the diffusive boundary layer surrounding phytoplankton cells^7^—and recent work has elucidated the diverse components of dissolved organic carbon (DOC), nitrogen (DON), and sulfur (DOS) involved in these exchanges^3^, together with the surprising taxonomic diversity and functional potential of associated bacteria (or the microbiome)^8^. Despite these recent advances, the signaling mechanisms that enable these taxa to recognize each other, regulate beneficial interactions, and facilitate metabolite exchanges are largely unknown. Here we identify for the first time an unrecognized role of vitamin B6 catabolites as communication molecules that mediate symbiotic interactions between the ubiquitous marine diatom *Asterionellopsis glacialis* and the planktonic Rhodobacteraceae *Pseudosulfitobacter pseudonitzschiae* and show the widespread importance of vitamin B6 in inter-microbial interactions in the ocean.

## Diatom DON Selectively Assimilated by Rhodobacteraceae

We have previously shown that *A. glacialis* selectively enriches members of the Rhodobacteraceae in its microbiome by secreting several DON metabolites as well as azelaic acid, which promotes the growth of Rhodobacteraceae while inhibiting opportunistic bacteria, such as Alteromonadaceae^9,10^. To examine the full potential of DON metabolites provided by the diatom to Rhodobacteraceae, we labeled axenic *A. glacialis* with ^15^N-nitrogen, then exchanged the medium and added three unlabeled bacteria: two beneficial Rhodobacteraceae, *P. pseudonitzschiae* F5 and *Phycobacter azelaicus* F10, and an opportunist, *Alteromonas macleodii* F12^9,11^. Using nanoSIMS, we examined ^15^N transfer from the diatom to the bacteria (Fig. 1a-c). After 12 hours of co-culture, only the Rhodobacteraceae assimilated 11 times more ^15^N-labeled DON than natural abundance levels. In comparison, *A. macleodii* had negligible ^15^N assimilation levels, which were indistinguishable from the background (Fig. 1c,d). This differential assimilation of DON is consistent with the diatom secretion of azelaic acid that inhibits *A. macleodii* growth for up to 24 hours^10^. We then quantified DON assimilation rates by Rhodobacteraceae via nanoSIMS using ^15^N-labeled *A. glacialis* co-cultured with *P. pseudonitzschiae* only. These measurements revealed that *P. pseudonitzschiae* rapidly assimilated ^15^N at a rate of 1.4±0.7 amol N cell^-1^ hr^-1^ (Fig. 1e,f), consistent with DON assimilation rates by marine bacteria in surface oceans^12^.

**Figure 1.**
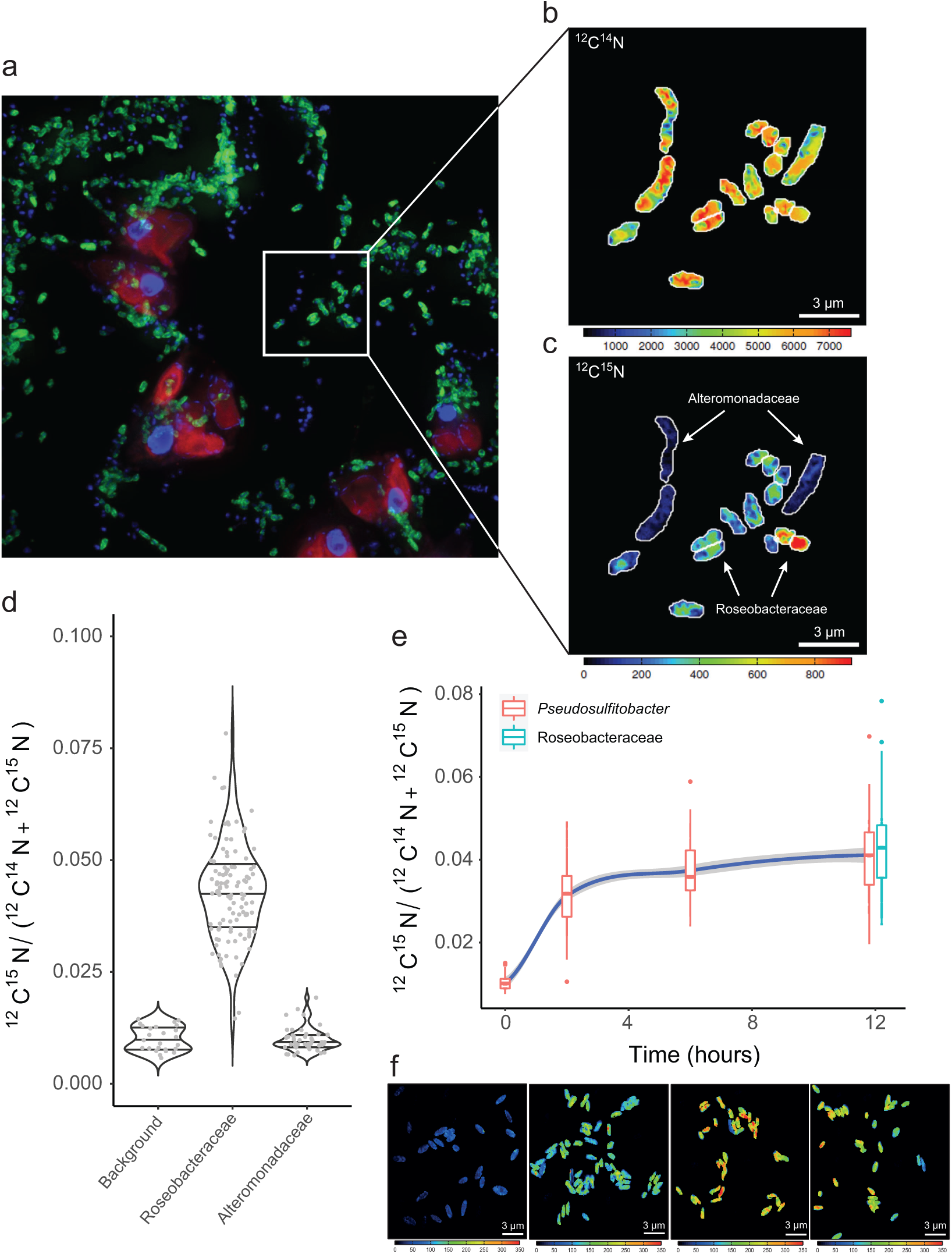
Characterization of diatom-derived DON uptake by beneficial bacteria by nanoSIMS. (a) CARD-FISH image of ^15^N-labeled *Asterionellopsis glacialis* co-cultured for 12 hours with *Alteromonas macleodii*, *Phycobacter azelaicus* and *Pseudosulfitobacter pseudonitzschiae*. The diatom nucleus is stained with DAPI (purple) and the plastid appears in red, while the roseobacters (*Phycobacter* and *Pseudonitzschiae*) are green and *A. macleodii* appears in blue. (b) Abundance of ^14^N (measured as ^12^C^14^N) in the window shown indicate ^14^N amounts across all cells. (c) Uptake of ^15^N (measured as ^12^C^15^N) in the window shown indicating differential uptake of ^15^N by roseobacters. (d) Ratio of ^15^N/^14^N uptake by bacteria across several windows showing only roseobacters accumulate diatom-derived ^15^N. (e and f) Time-course of *P. pseudonitzschiae* uptake of diatom-derived ^15^N. Values for Roseobacteraceae represent uptake at 12 hours by both *P. pseudonitzschiae* and *P. azelaicus* and were extracted from (d).

## Characterizing the Diatom DON and uptake by bacteria

To decipher the chemical complexity of the secreted diatom DON, ^15^N-DON was extracted and concentrated from the *A. glacialis* supernatant using solid-phase extraction. To aid in annotating ^15^N-containing metabolites, two commercial chemical libraries (containing 343 nitrogen-containing natural abundance metabolites at defined concentrations and a ^13^C-labeled yeast extract as internal standard) (Supplementary Information Table 1) were spiked in the extract and the mixture was analyzed on a Bruker Fourier-transform ion cyclotron resonance mass spectrometer (FT-ICR-MS). Fragmentation patterns (tandem MS) of metabolites containing ^15^N-atoms were compared to those from the natural abundance metabolite library, enabling the putative annotation of diatom metabolites (Extended Data Fig. 1a). Using this method, 208 diatom metabolites were putatively annotated that belonged mostly to amino acids and their derivatives, nucleosides and nucleotides (Extended Data Fig. 1b, Supplementary Information Tables 2 and 3).

To determine which of these putative DON metabolites are taken up by Rhodobacteraceae, *P. pseudonitzschiae* cells were incubated for 3 hours in cell-free ^15^N-labeled *A. glacialis* spent medium, after which bacteria were removed by filtration to obtain a second spent medium. ^15^N-labeled *A. glacialis* was reintroduced to the second spent medium and incubated for 3 additional hours, followed by filtration to obtain a third spent medium. All three spent media were extracted and analyzed with FT-ICR-MS (Fig. 2a). A comparison between the relative abundance of ^15^N-labeled putative metabolites revealed that within 3 hours, *P. pseudonitzschiae* took up 53 metabolites simultaneously, most of which belonged to amino acids and derivatives, benzenoids, and indoles and derivatives and representing 25% of putatively annotated diatom ^15^N-metabolites (Fig. 2b, Extended Data Figs. 1b and 2, Supplementary Information Table 4). Putative diatom metabolites taken up by *P. pseudonitzschiae* had on average statistically lower molecular weight and higher hydrophobicity than those not taken up by the bacterium (Extended Data Fig. 1c-e), which is consistent with metabolites that are less energetic to transport across cell membranes^13^. Surprisingly, *P. pseudonitzschiae* secreted two putative ^15^N-metabolites, 4-pyridoxate (4-PA) and phosphoethanolamine (PEtN), from diatom ^15^N-labeled precursors. Indeed, the bacterium took up diatom-derived 4-pyridoxolactone, pyridoxamine, and pyridoxal, all of which are related to vitamin B6 and may have served as precursors for 4-PA biosynthesis by the bacterium (Fig. 2b, Supplementary Information Table 4). While pyridoxal-phosphate (PLP) and, to a lesser extent, pyridoxamine-phosphate (PMP) are considered the active forms of vitamin B6^14^, pyridoxamine, pyridoxal, 4-pyridoxolactone and 4-PA are considered intermediates or catabolites of PLP and PMP that do not carry out any biological or ecological functions^4,15^. Indeed, most bacteria excrete 4-PA as a non-salvageable, waste product^16^. PEtN is derived from ethanolamine and is a precursor of phospholipids, especially phosphatidylethanolamine, a major membrane lipid across the tree of life^17^. A comparison between the relative abundance of ^15^N-labeled putative metabolites in the third vs the second spent medium indicates that *A. glacialis* replenished the metabolites taken up by the bacterium, and importantly, took up ^15^N-labeled 4-PA and PEtN, suggesting a communication mechanism between both taxa (Fig. 2b, Supplementary Information Table 4). These findings suggest that for the first time these seemingly ecologically inert molecules with no prior role in interspecies interactions may be acting as signals between the diatom and its bacterial partner.

**Figure 2.**
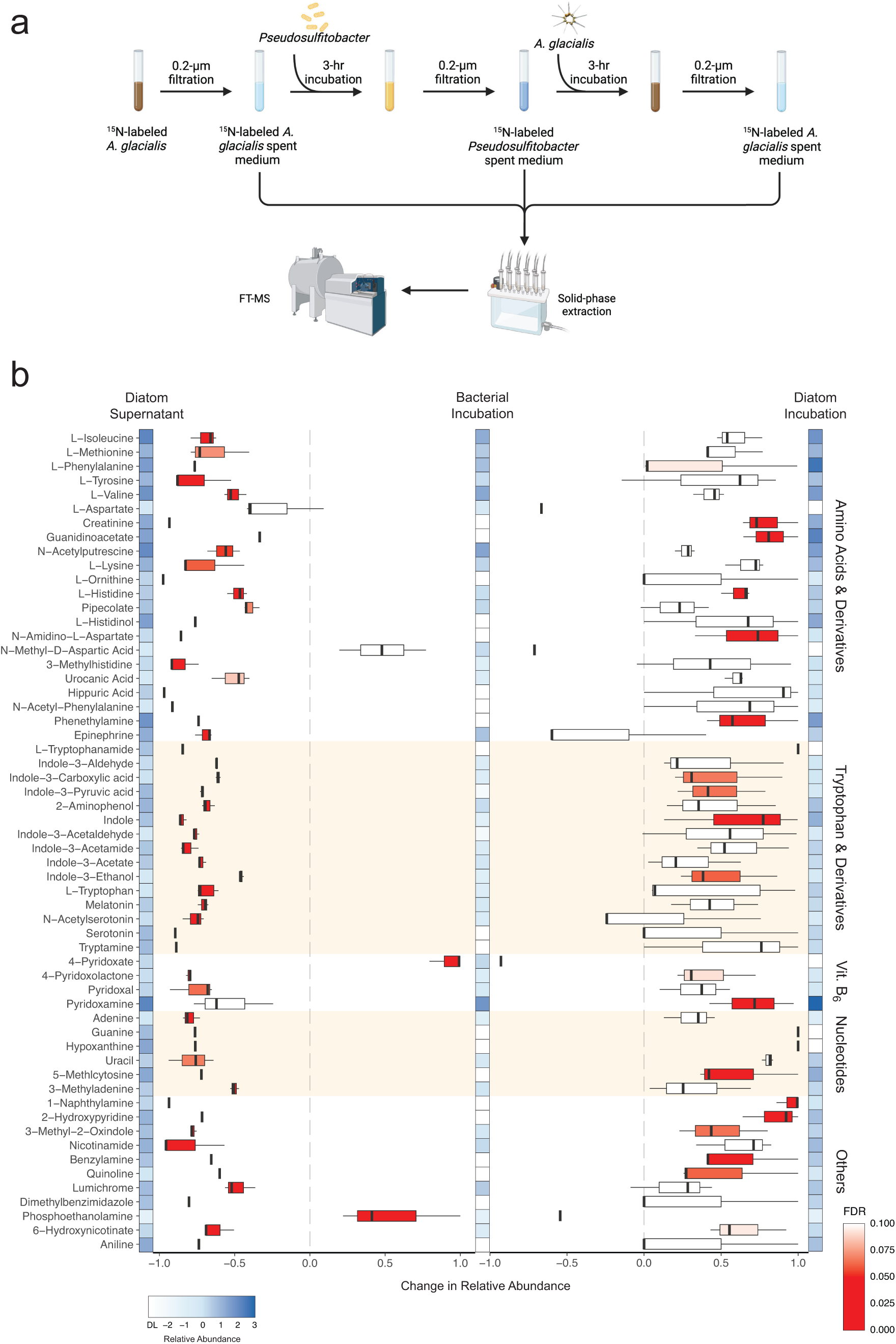
Uptake of diatom-derived ^15^N metabolites by *P. pseudonitzschiae*. (a) Scheme for the incubation experiments. (b) Comparison between (left) the bacterial incubation in diatom spent media (2^nd^ spent medium in (a)) vs the original diatom spent medium and between (right) the reintroduced diatom spent medium (3^rd^ spent medium in (a)) vs the bacterial incubation. Fragmentation patterns of all putative ^15^N-metabolites vs natural abundant metabolites are shown in Extended Data Fig. 2. Heat map represents a rough approximation of the concentration of metabolites based on a 1-point calibration using internal standards for each metabolite. Change in relative abundance is shown, where negative values indicate bacterial (left) or diatom (right) uptake and positive values indicate production. Spent media control were incubated at the same conditions for each step to account for abiotic degradation. All relative abundances were calculated based on n=5 samples.

## Deciphering signaling between the diatom and bacterium

To examine whether pyridoxal, pyridoxamine, 4-pyridoxolactone, 4-PA and PEtN are involved in a signaling mechanism between both taxa, we grew each partner separately in defined media supplemented with only one of these metabolites and conducted RNA-seq to assess transcriptional responses to these putative signals at 60 and 180 minutes post-addition (Extended Data Table 1). Addition of 4-PA or PEtN to the diatom induced a moderate transcriptional response, yielding an average of 567 differentially expressed genes (DEGs) across both time points (log2FC>±1, *p*adj<0.05; 0.3-5% of all diatom CDS) (Extended Data Table 1, Supplementary Discussion, Supplementary Information Tables 14-17). Likewise, addition of pyridoxamine, 4-pyridoxolactone, or pyridoxal induced a relatively small to moderate transcriptional response in the bacterium, yielding an average of 82 DEGs across both time points (0.08-5.3% of all bacterial CDS) (Extended Data Table 1, Supplementary Discussion, Supplementary Information Tables 6-11). Addition of pyridoxine to the bacterium, a putative diatom-derived vitamin B6 intermediate not taken up (Supplementary Information Table 3), induced no transcriptional response by the bacterium, indicating the bacterial responses to the other vitamin B6 intermediates are specific (Extended Data Table 1, Supplementary Information Table 5).

Since the diatom and bacterium likely encounter these signal molecules simultaneously, we repeated the RNA-seq experiments in the presence of both 4-PA and PEtN for the diatom and pyridoxamine, 4-pyridoxolactone, and pyridoxal for the bacterium. Surprisingly, addition of 4-PA and PEtN to the diatom led to an increase in differential expression to 1893 and 3293 genes at 60 and 180 mins, respectively, compared to 788 and 728 at 60 and 180 mins for 4-PA alone or 710 and 44 at 60 and 180 mins for PEtN alone (Extended Data Table 1). Indeed, the transcriptional response of the diatom to both signals yielded 3274 unique DEGs that were not differentially expressed with either signal alone (Extended Data Fig. 3a), suggesting a synergistic effect that can only be observed when both signals are co-perceived by the diatom. When *P. pseudonitzschiae* was exposed to 4-pyridoxolactone, pyridoxamine and pyridoxal simultaneously, it differentially expressed 24 and 897 genes at 60 and 180 mins, respectively, compared to a maximum of 257 DEGs observed with pyridoxal (Extended Data Table 1). Like the diatom, the bacterial transcriptional response to the combination of all three signals yielded 694 unique DEGs that were not differentially expressed with any of the other signals alone (Extended Data Fig. 3b), suggesting again a synergistic effect for these signals. Curiously, simultaneous exposure to this mixture of signaling molecules led to a differential expression of ∼20% of all CDS in both the diatom and bacterial genomes. Despite the broad effect of these signals on the transcriptional profiles of the diatom and bacteria, these molecules had no discernible influence on the growth of either organism.

## Transcriptional patterns induced by signals

Upon 60 minutes of exposure to 4-PA and PEtN, the diatom underwent major restructuring of metabolic pathways that indicate repression of growth and biosynthesis. This included a significant downregulation in several pathways (padj<0.05), including carbon fixation, fatty acid biosynthesis and metabolism and the ribosome (Extended Data Fig. 4). At 180 minutes, most of these pathways remained downregulated; however, the diatom transcriptome showed signs of nuclear and growth reactivation and redox regulation. Enriched upregulated pathways included the cell cycle, homologous recombination, nucleotide biosynthesis, DNA replication, pentose phosphate, and glutathione metabolism. These broad patterns suggest that the diatom initially suppresses autotrophic growth to conserve resources after encountering the bacterial signals, followed by major reprogramming to induce cell division and metabolic recycling to enter a symbiotic state, which should lead to increased carbon flow from the diatom to the bacterium. Similar patterns of repressing carbon fixation and autotrophic growth followed by metabolic reprogramming have been documented for photosynthetic organisms, such as lichen-like associations, plants and other phytoplankton, after exposure to symbiotic partners^18–22^. In contrast, *P. pseudonitzschiae* only exhibited enriched pathways after 180 minutes of exposure to 4-pyridoxolactone, pyridoxamine and pyridoxal that all were downregulated, including quorum sensing, chemotaxis, nitrogen metabolism, secretion systems and transporters (Extended Data Fig. 4). These patterns may suggest preparation for phycosphere colonization and mechanisms initiating attachment.

A closer examination of the *A. glacialis* response to 4-PA and PEtN indicates upregulation of the biosynthesis genes of many amino acids and other metabolites shared with the bacterium. These include methionine, leucine, isoleucine, valine, aspartate, glutamine (via urea degradation), glutamate, tyrosine, phenylalanine and tryptophan (Fig. 3, Supplementary Information Tables 18-19). The bacterial signals further induced the diatom to convert tryptophan to (i) indole-3-acetate (IAA) via tryptamine and indole-3-acetaldehyde, and (ii) serotonin, acetyl-serotonin and melatonin, all of which were shared with the bacterium (Fig. 2b, Fig. 3). These patterns suggest the bacterial signals prime the diatom to prepare for its mutualistic partner by increasing the biosynthesis of a variety of metabolites, while potentially anticipating the production of bacterial-derived urea that can then be used to synthesize glutamine.

**Figure 3.**
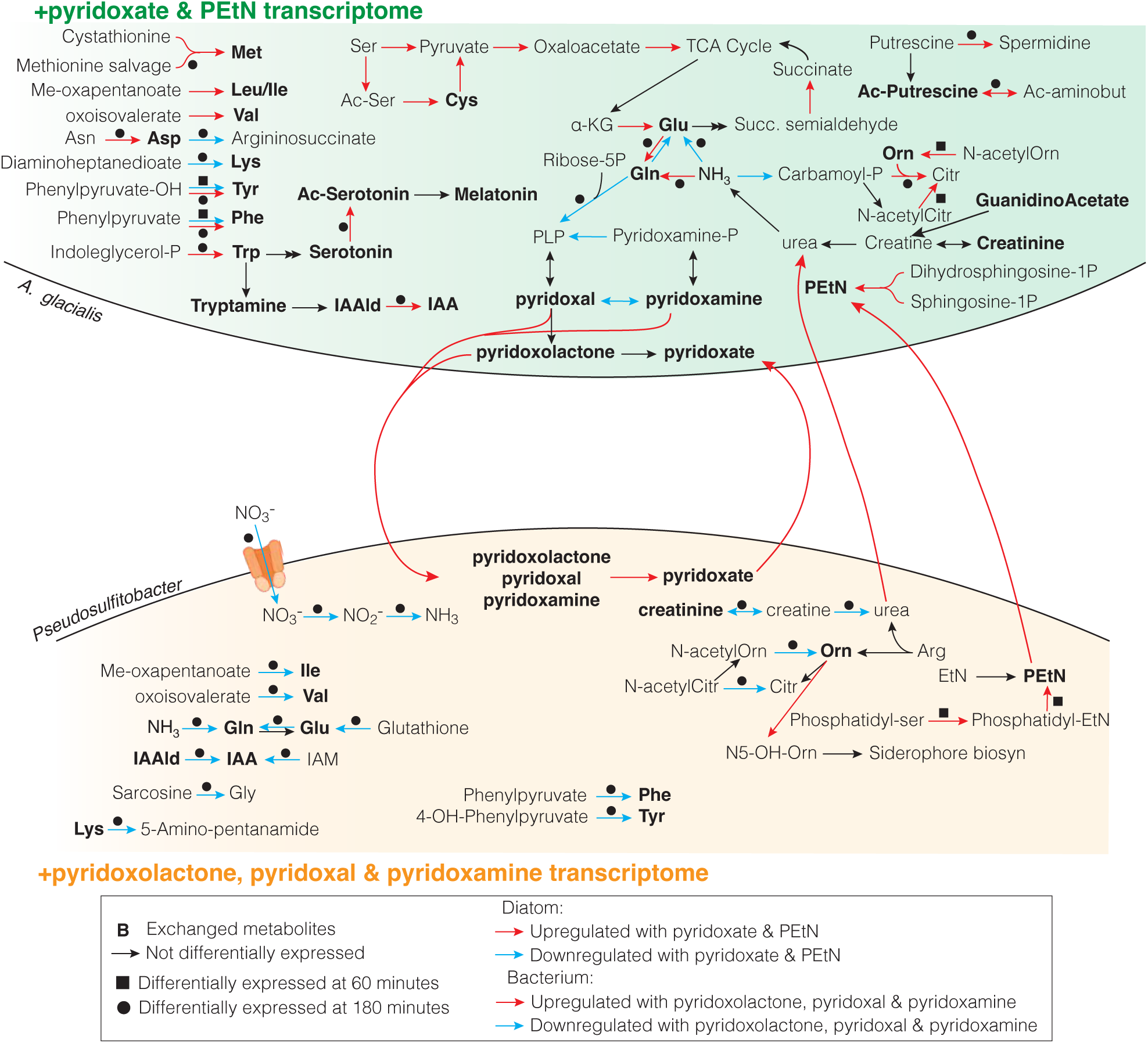
Transcriptional patterns in metabolism in *A. glacialis* (top) and *P. pseudonitzschiae* (bottom) in response to pyridoxate+PEtN and 4-pyridoxolactone+pyridoxal+pyridoxamine, respectively. Genes were considered differentially expressed with the following cutoffs based on biological triplicates (log_2_fc ≥ ±1.0 p_adj_ < 0.05). Boldfaced metabolite names/abbreviations are exchanged in Fig. 2b. Upregulated genes are represented by red arrows, downregulated genes by cyan arrows, and not differentially expressed genes by black arrows. Squares and circles indicate differential expression at 60 or 180 minutes, respectively. All transcriptomes were based on biological triplicates. List of abbreviations: all amino acids are represented by the standard 3-letter abbreviations for the 20-essential amino acids; 4-hydroxyphenylpyruvate; Indole-glycerol-P=indole-glycerol phosphate; Ac-serotonin=acetyl-serotonin; IAA=indole-3-acetate; IAAld=indole-3-acetaldehyde; IAM=indole-3-acetamide; Ac-ser=acetyl-serine; PLP=pyridoxal phosphate; Pyridoxamine-P=pyridoxamine phosphate; α-KG=alpha-ketoglutarate; Succ. Semialdehyde=succinate semialdehyde; Ribose-5P=ribose-5-phosphate; Carbamoyl-P=carbamoyl phosphate; Orn=ornithine; Ac-putrescine=acetyl-putrescine; Ac-aminobut=Acetyl-aminobutyric acid (GABA); Citr=citrulline; Sphingosine-1P=sphingosine-1-phosphate; Me-oxapentanoate=methyl oxopentanoate; N5-OH-Orn=N5-hydroxy ornithine.

In response to 4-pyridoxolactone, pyridoxamine and pyridoxal, *P. pseudonitzschiae* differentially regulated genes involved in the biosynthesis and catabolism of some of the 53 diatom-derived metabolites taken up by the bacterium, mainly at 180 mins. Biosynthesis of many amino acids and derivatives secreted by the diatom and taken up by the bacterium were downregulated, including phenylalanine, tyrosine, glutamate, glutamine, isoleucine, valine, and IAA (Fig. 3, Supplementary Information Tables 12-13). Uptake and reduction of nitrate to ammonia were downregulated, reflecting the broad pattern of uptake of diatom-derived DON. Ornithine, also secreted by the diatom, was converted to N5-hydroxyornithine, a building block for siderophore backbones^23^. Siderophores are critical for bacterial iron acquisition, and can enable diatoms and other phytoplankton to acquire otherwise inaccessible iron^24^. Biosynthesis of PEtN was upregulated from phosphatidyl-ethanolamine at 60 mins, suggesting that the diatom signals induce the production of PEtN. Interestingly, similar patterns of expression related to diatom-derived metabolites were also observed in the bacterium in the presence of 4-pyridoxolactone, pyridoxamine, or pyridoxal added separately (Supplementary Discussion, Supplementary Information Tables 6-11). For example, the biosynthesis of valine, isoleucine, methionine, PEtN and creatinine were upregulated. Creatinine was further converted to creatine and then to urea in the presence of pyridoxolactone or pyridoxal. Because urea appears central to the diatom transcriptional response, we examined the ability of *P. pseudonitzschiae* to secrete urea. While *P. pseudonitzschiae* controls grown in regular media produced 25 μM urea, cells grown with 4-pyridoxolactone showed a 3-fold increase in urea production (Extended Data Fig. 5). In conclusion, these patterns suggest that the diatom signals prime the bacterium to anticipate the uptake of diatom DON metabolites and to secrete specific metabolites in response, mainly urea as a nitrogen source and PEtN and 4-PA as signals.

## Characterizing the proteins synthesizing pyridoxolactone and 4-pyridoxate

Because vitamin B6 is essential for all life, we examined the genetic potential of the diatom and bacterium to produce vitamin B6, as well as the four B6-related signals identified. Mining the diatom and bacterial genomes indicated that both are capable of converting pyridoxamine phosphate to the active form of the vitamin, pyridoxal phosphate (PLP) (Extended Data Fig. 6). In addition, the diatom is capable of the conversion between PLP and pyridoxal, and pyridoxamine phosphate and pyridoxamine. However, no clear homologs were identified in the diatom genome for pyridoxal 4-dehydrogenase (PLDH), which converts pyridoxal to 4-pyridoxolactone, nor were homologs for pyridoxal oxidase and/or pyridoxolactonase (PDLA) that convert either pyridoxal or 4-pyridoxolactone, respectively, to 4-PA in the bacterial genome (Extended Data Fig. 6). PLDH belongs to the NAD-dependent short-chain dehydrogenase family^25^; therefore, we identified 15 putative proteins in the diatom genome belonging to this family without metabolite-specific annotations (Supplementary Information Table 20). These 15 proteins were modeled with AlphaFold^26^ to generate tertiary protein structures to compare with the existing crystal structure of PLDH from *Mesorhizobium loti*^25^. This analysis yielded two potential protein homologs in *A. glacialis* with percent amino acid identities >20%, mostly conserved active site residues, and RMSD <2.2. These two homologs were then expressed in *Escherichia coli* and purified (Supplementary Information Table 22). To test the activities of these proteins *in vitro*, we measured the production of 4-pyridoxolactone when pyridoxal was incubated alone, when PLDH homologs were incubated alone, or when PLDH homologs were incubated with pyridoxal. Only one homolog (A3_.00g105710.m01) led to the production of 4-pyridoxolactone and complete consumption of the substrate within 60 minutes of incubation (Fig. 4a-c, Supplementary Video 1, Supplementary Data 1).

**Figure 4.**
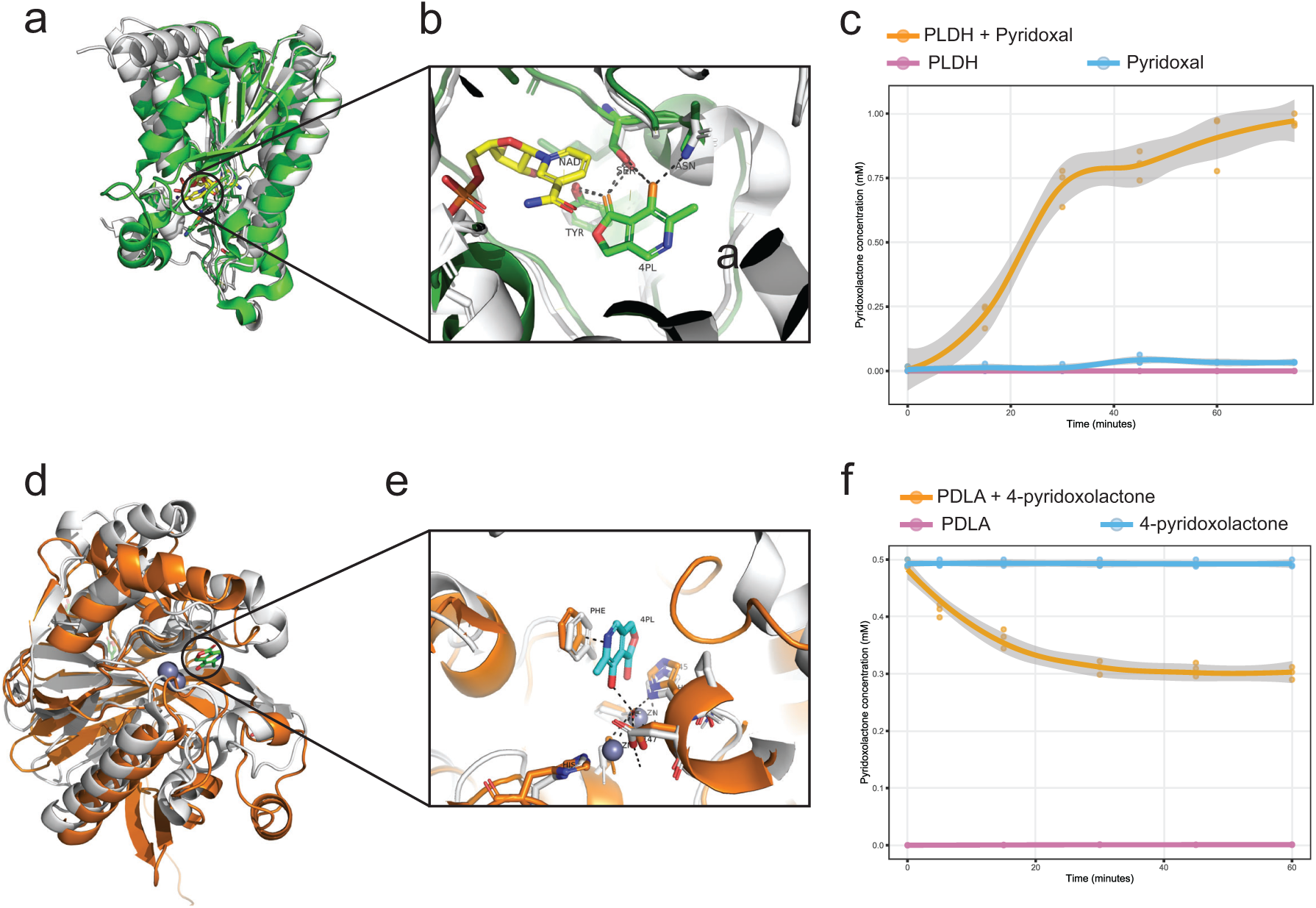
Enzyme structure and activity for PLDH and PDLA from *A. glacialis* and *P. pseudonitzschiae,* respectively. (a) Alignment of the folded *A. glacialis* PLDH homolog (green) with 3RWB (white) imported from the Protein Database (PDB) from *Mesorhizobium loti*. (b) View of the aligned active site of both proteins, showing 4-pyridoxolactone and NAD+. (c) *In vitro* enzyme activity of *A. glacialis* PLDH showing production of 4-pyridoxolactone. (d) Alignment of the folded *P. pseudonitzschiae* PDLA homolog (brown) with 4KEQ (white) imported from PDB from *Mesorhizobium japonicum*. (e) View of the aligned active site of both proteins, showing 4-pyridoxolactone. (f) *In vitro* enzyme activity of *P. pseudonitzschiae* PDLA showing consumption of 4-pyridoxolactone.

Mining the bacterial genome indicated the presence of two potential PDLA homologs that belonged to proteins containing metallo-beta-lactamase/lactonase domains^27^ (Supplementary Information Table 21). As above, we used AlphaFold^26^ to generate tertiary structures of each protein sequence and compared them to the existing crystal structure of PDLA from *M. japonicum*^15^. This analysis yielded one potential protein homolog in *P. pseudonitzschiae* (IHQ53_RS05450) with percent amino acid identity >20%, conserved active site residues, and RMSD ≤3.0. This protein was expressed in *E. coli* and we measured its activity *in vitro* as described above (Supplementary Information Table 22). We measured the consumption of 4-pyridoxolactone when it was incubated alone, when the PDLA homolog was incubated alone or when both were incubated together. The presence of the PDLA homolog led to the consumption of 40% of the substrate within 30 mins of incubation, after which protein activity was halted likely due to instability in solution (Fig. 4d-f, Supplementary Video 2, Supplementary Data 2). These findings indicate that the diatom and bacterium are capable of biosynthesizing 4-pyridoxolactone and 4-pyridoxate, respectively, via new homologs of PLDH and PDLA.

## Mapping the abundance of PLDH and PDLA in public genomes and in the Oceans

The detection of genes and transcripts coding for PLDH and PDLA can serve as a proxy for vitamin B6-mediated communication among eukaryotes and prokaryotes, because these genes synthesize catabolites that are not precursors to vitamin B6 biosynthesis (i.e., 4-pyridoxolactone), or are excreted, waste products (i.e., 4-PA)^4,15^. Using the known homologs of PLDH from *A. glacialis*, *M. loti* and *M. japonicum*, and PDLA from *P. pseudonitzschiae* and *M. japonicum*, we constructed HMM profiles for each protein and mined publicly available genomes (RefSeq) for their taxonomic distribution across the tree of life. PLDH putative homologs were widely distributed across the eukaryotic domain, especially in photosynthetic organisms, including plants (Streptophyta) and phytoplankton (Bacillariophyta and Chlorophyta), and to a lesser extent in prokaryotes (mostly Bacteriodota) (Extended Data Fig. 7a, Supplementary Information Table 23). PDLA putative homologs were widely distributed across bacterial phyla with the majority belonging to Pseudomonadata (alpha-, beta- and gammaproteobacteria) and Actinomycetota (Extended Data Fig. 7b, Supplementary Information Table 24). No homologs were found in the eukaryotic or archaeal domains, indicating only bacteria are capable of producing 4-PA. These findings suggest that this communication mechanism is broadly distributed across eukaryotic-prokaryotic taxa.

Using the HMM profiles for PLDH and PDLA, we also mined the Tara Oceans database for the abundance and activity of these proteins, with the requirement that both genes or transcripts must be present or expressed simultaneously at the same depth and station to be indicative of potential signaling. Using these criteria, both genes were present in the Tara Oceans database in 78 distinct samples (75% of total samples investigated) across the surface and the deep chlorophyll maximum (Fig. 5a, Extended Data Fig. 8a). Eukaryotic taxa that contributed the most to *pldh* homologs were dominated by diatoms (Bacillariophycidae, Thalassiosiophycidae, Fragilariophycidae, Cymatosirophycidae, Chaeotocerotophycidae, Corethrophycidae), Embryophyta, Bilateria and several phytoplankton groups, such as green algae (Mamiellophyceae), ochrophytes (*Aureococcus*), prymnesiophytes (*Phaeocystis*), green algae (Mamiellophyceae), coccolithophores (*Emiliania*) and dinoflagellates (Symbiodiniaceae). Prokaryotic taxa that contributed the most to *pdla* were by far dominated by roseobacters (Rhodobacteraceae) in addition to other families including Pseudomonadaceae, Bradyrhizobiaceae, Burkholderiaceae, Phyllobacteriaceae and Alcaligenaceae (Fig. 5a, Extended Data Fig. 8a).

**Figure 5.**
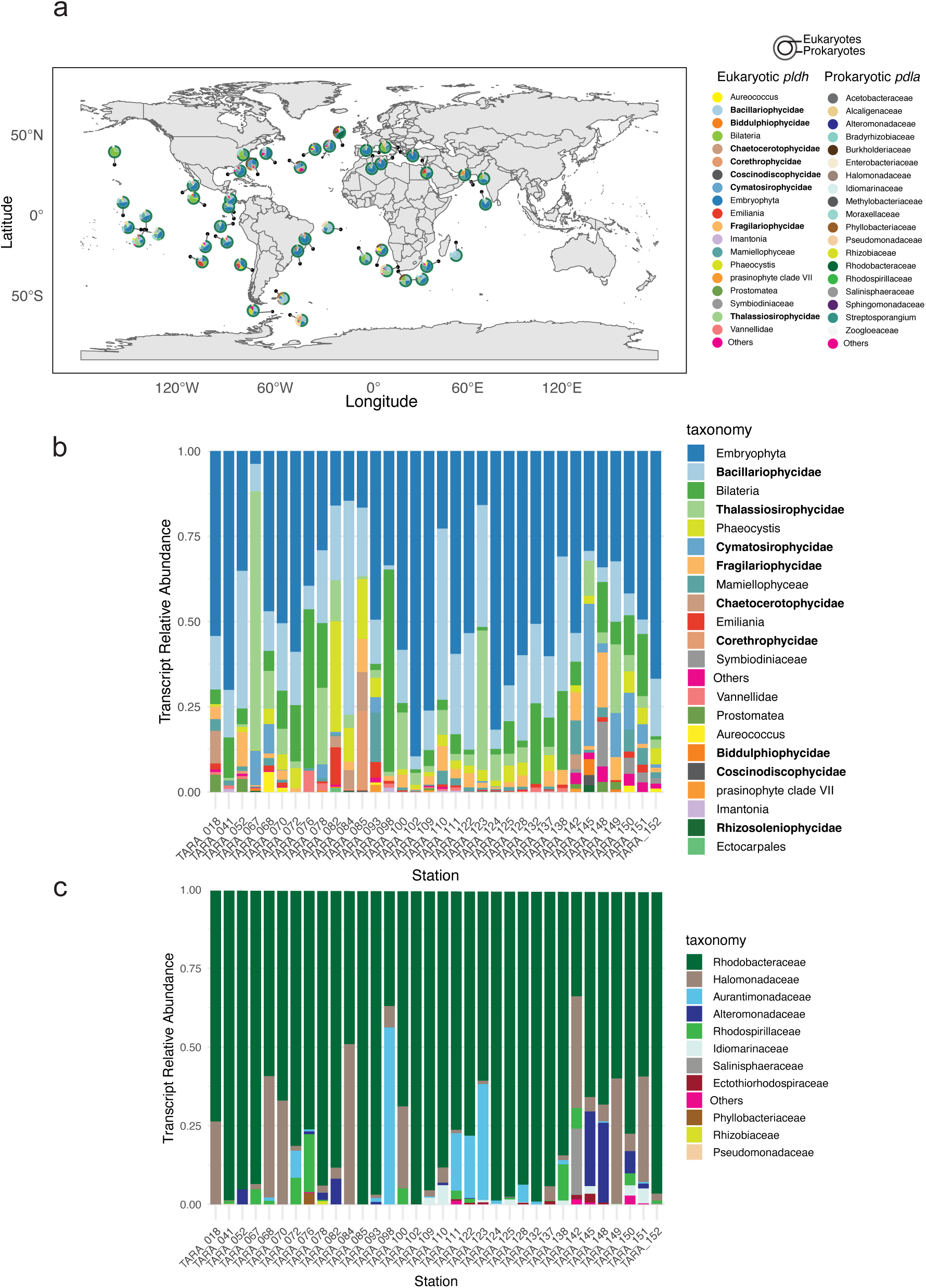
Abundance of *pldh* and *pdla* genes and transcripts in the surface ocean. HMM profile of each protein sequence was used to mine the Tara Oceans database surface samples across all available size fractions. (a) Relative abundance and co-localization of *pldh* and *pdla* in Tara metagenomes in the surface. (b) Transcript relative abundance of *pldh* in Tara metatranscriptomes in the surface. All bold-face taxa are diatoms. (c) Transcript relative abundance of *pdla* in the same stations as (b) in Tara metatranscriptomes in the surface.

In addition, we mined the Tara Oceans metatranscriptome data and used the same criteria used for metagenomic reads. Transcripts from both genes were co-expressed at 67 distinct samples (64.4% of total samples investigated) across the surface and the deep chlorophyll maximum (Fig. 5b,c, Extended Data Fig. 8b,c). Similar to metagenomes, relative transcript abundance for *pldh* were dominated by diatoms, other phytoplankton groups as well as embryophyta and bilateria. Relative transcript abundance for *pdla* were dominated again by roseobacters with also a significant contribution from Halomonadaceae, Aurantimonadaceae and Rhodospirillaceae. These findings indicate that eukaryotic phototroph-prokaryotic heterotroph interactions may be mediated by these enzymes and the molecules they synthesize.

## Discussion

The discovery of vitamin byproducts acting as communication signals between eukaryotes and prokaryotes constitutes a substantial conceptual shift in our understanding of how chemical communication structures interkingdom interactions. B-vitamins (e.g., B1, B3, B6, B7 and B12) are classically thought to be cofactors for enzymes^28^ and significant attention has been devoted to exchanges and cross-feeding among marine microbes of some of these vitamins, especially vitamin B12^29^, which plays a critical role in methionine biosynthesis and methylmalonyl-CoA metabolism^30^. Vitamin B6 serves as a cofactor for more than 140 enzymes that are involved in the biosynthesis of amino acids, heme, neurotransmitters, vitamin B3, and the metabolism of homocysteine among others^31^. Because of its broad requirement in metabolism^32^, only a few microbes are known to be auxotrophs for vitamin B6 and thus exchange of vitamin B6 among microbes is uncommon^33^. We have shown for the first time that vitamin B6 intermediates and catabolites—pyridoxal, pyridoxamine, 4-pyridoxolactone and 4-PA—are not ecologically inert but, together with PEtN, act as signals that initiate metabolite exchange between the diatom and its bacterial partner.

4-PA is widely considered an irreversible waste product of vitamin B6 metabolism^34^, which is further degraded by *Pseudomonas putida*, *Arthrobacter* spp., and *Nocardia* spp. to succinate^14,35^. Most other taxa, such as many enteric, soil and marine bacteria, cannot metabolize 4-PA; as a result, 4-PA accumulates as a terminal product and is excreted^36^. Here, we show for the first time that this waste product is in fact an interkingdom signaling molecule. We revealed the ubiquity of genes and transcripts of eukaryotic *pldh* and prokaryotic *pdla* in RefSeq and Tara Oceans as proxy for such signaling between the eukaryotic and prokaryotic domains in the oceans and beyond. While we recognize that the presence/expression of *pldh* and *pdla* does not prove active signaling between eukaryotic hosts and bacteria *a priori*, the ubiquity of these genes and their active co-expression among phototrophs and heterotrophs in the oceans suggest a widely used communication. Specifically, the broad presence of 4-PA excretion across the bacterial domain, irrespective of environment, suggests that it may have an important and overlooked evolutionary role in regulating interactions with eukaryotic hosts, particularly photosynthetic ones. This is further corroborated by the broad distribution of *pldh* in plant genomes (Extended Data Fig. 7).

The synergistic effect of multiple communication molecules on gene regulation, as observed here, has been reported in only a few other biological and symbiotic systems. For example, when *Pseudomonas aeruginosa* is exposed to the quorum sensing signals 3-oxo-C12 and C4-homoserine lactones, it differentially regulates a greater number of secretome genes than the sum of each signal alone^37,38^. During nodulation in legumes, ethylene and jasmonic acid both act combinatorially to regulate nodulation in the plant and Nod factor signal transduction in bacteria^39^. In the context of the phycosphere, the use of multiple signals by phytoplankton and bacteria to coordinate interactions with their partner may be a strategy to minimize or eliminate cheaters^40^. Yet, the broad taxonomic distribution of this communication mechanism seems incompatible with an arms race between bacterial symbionts and cheaters. Indeed, cheaters may easily establish a foothold in the phycosphere via horizontal gene transfer of such widely distributed communication genes. However, eukaryotic-prokaryotic interactions seldom involve a single or even a pair of signals. For example, legume-rhizobia symbiosis utilizes a cascade of signals from each partner that ensures symbiosis is established with the right partner with strain-level specificity^41^. Moreover, some canonical signals in this symbiosis, such as IAA, are broadly present across many symbiotic bacterial groups (e.g., *Bradyrhizobium*, *Mesorhizobium*, *Burkholderia*) as well as pathogens (e.g., *Pseudomonas*)^42^. These patterns suggest that a cascade of signals is required to ensure the integrity of symbiotic interactions. Likewise, vitamin B6 catabolites and PEtN are likely some of many signals that regulate metabolite exchanges to ensure that cheaters cannot find their way into the phycosphere.

## Conclusion

Our work reveals that vitamin B6 catabolites—long regarded as mere metabolic intermediates or waste products—serve as potent, bidirectional chemical signals that structure interactions between a diatom and its bacterial partner. By demonstrating that these molecules elicit coordinated, combinatorial transcriptional reprogramming without altering growth, we identify a previously unrecognized layer of metabolic communication in the phycosphere. The discovery of dedicated enzymes that generate these signals, together with their broad distribution and expression across genomes, global ocean metagenomes and metatranscriptomes, suggests that vitamin B6–mediated signaling is a widespread organizing principle in marine microbial ecosystems and beyond. These findings expand the conceptual framework of nutrient exchanges in the sunlit portion of the oceans to include combinatorial chemical dialogue and raise the prospect that other metabolites canonically viewed as inert or degradative end products may act as information-rich signals underpinning interkingdom interactions in the world’s oceans.

## Supporting information

SI Discussion

## Acknowledgements

We would like to acknowledge the Gordon and Betty Moore Foundation’s (GBMF) award to S.A.A., J.B. Raina and J.R. Seymour, and Tamkeen awards under the NYU Abu Dhabi Research Institute Award to the NYUAD Center for Genomics and Systems Biology (ADHPG-CGSB) and to S.A.A. (AD179) for funding this work. We also would like to acknowledge the Mass Spectrometry Facility and the Core Technology Platform at NYU Abu Dhabi.

## Author Contribution

SAA, MAO, JBR and JRS conceived the project; MAO and SAA designed the experiments; MAO, LSYC, CF, SL and DT conducted experiments; MAO, ARM, SAA, SL and DT analyzed data; SAA, JBR and JRS acquired funding; all co-authors contributed to writing the manuscript.

## Data Availability

All mass spectrometry data are available in MassIVE under accession number MSV000100465. Diatom and bacterial genomes were previously deposited in GenBank under accession numbers GCA_014885115.2 and GCF_014805345.1, respectively. Diatom genome annotations and gene models was deposited on Zenodo (https://zenodo.org/records/18194605). All RNA-seq data are available in the short-read archive under accession numbers PRJNA1395043.

---

**Extended Data Figure 1.**
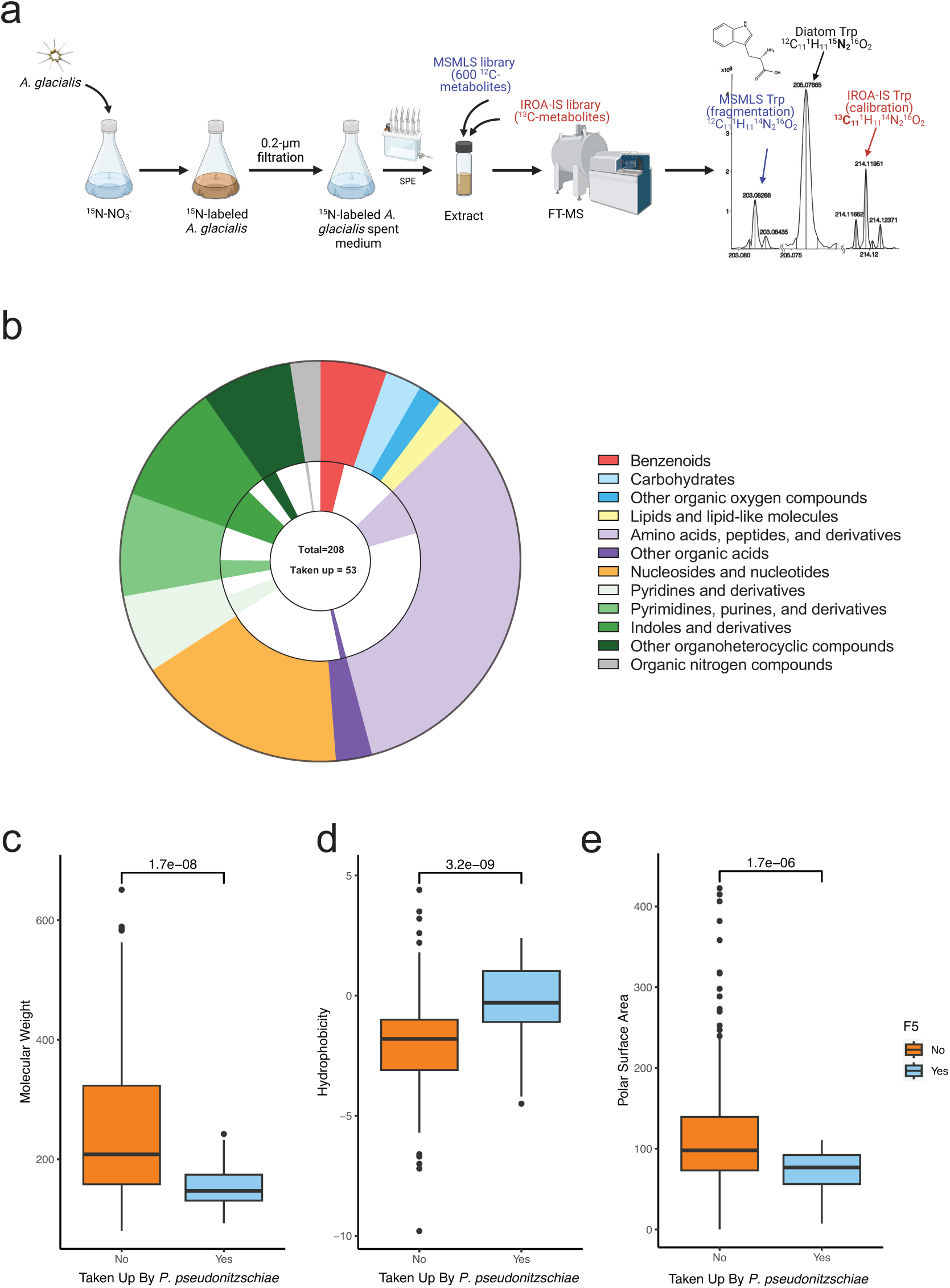
Characterization of the diatom dissolved organic nitrogen (DON) and its assimilation by Rhodobacteraceae. (a) Scheme for annotation of diatom DON excreted metabolites. *A. glacialis* was ^15^N-labeled by growth in ^15^N-NO_3-_, then cells were filtered to obtain a ^15^N-enriched spent medium and extracted using solid-phase extraction. Extracts were spiked with the MSMLS chemical library, comprised of 600 metabolites at natural abundance, and the IROA-IS chemical library, comprised of an extract of ^13^C metabolites. Signal response from the IROA-IS metabolites were used as internal standards to normalize signal across replicates, while the MSMLS were used for fragmentation to identify diatom metabolites. Samples were analyzed with FTMS. An example of peaks from tryptophan is shown. (b) Categories of putatively annotated ^15^N-metabolites from the diatom and those taken up by *P. pseudonitzschiae*. (c) Molecular weight profile of metabolites taken up or not taken up by *P. pseudonitzschiae*. (d, e) Polarity profile, examined through hydrophobicity and polar surface area, of metabolites taken up or not taken up by *P. pseudonitzschiae*. Statistical significance is indicated above each graph using a Student’s t-test.

**Extended Data Figure 2.**
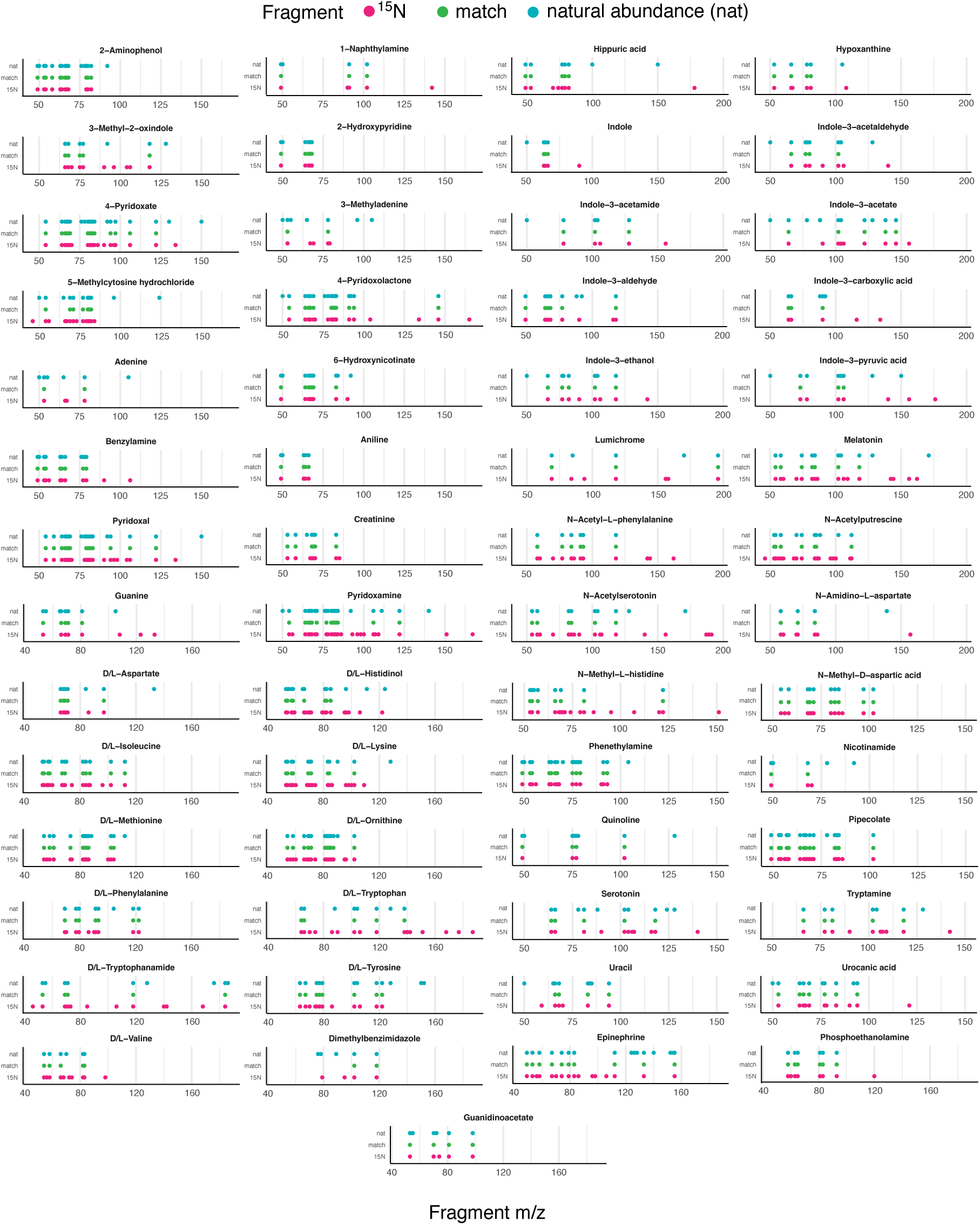
Fragmentation patterns of all putative metabolites displayed in Figure 2. Fragments are indicated by dots for natural abundant metabolites from the IROA library (cyan) and ^15^N metabolites from the diatom (red). If a fragment is matched between the two datasets, a matched dot is indicated (green).

**Extended Data Figure 3.**
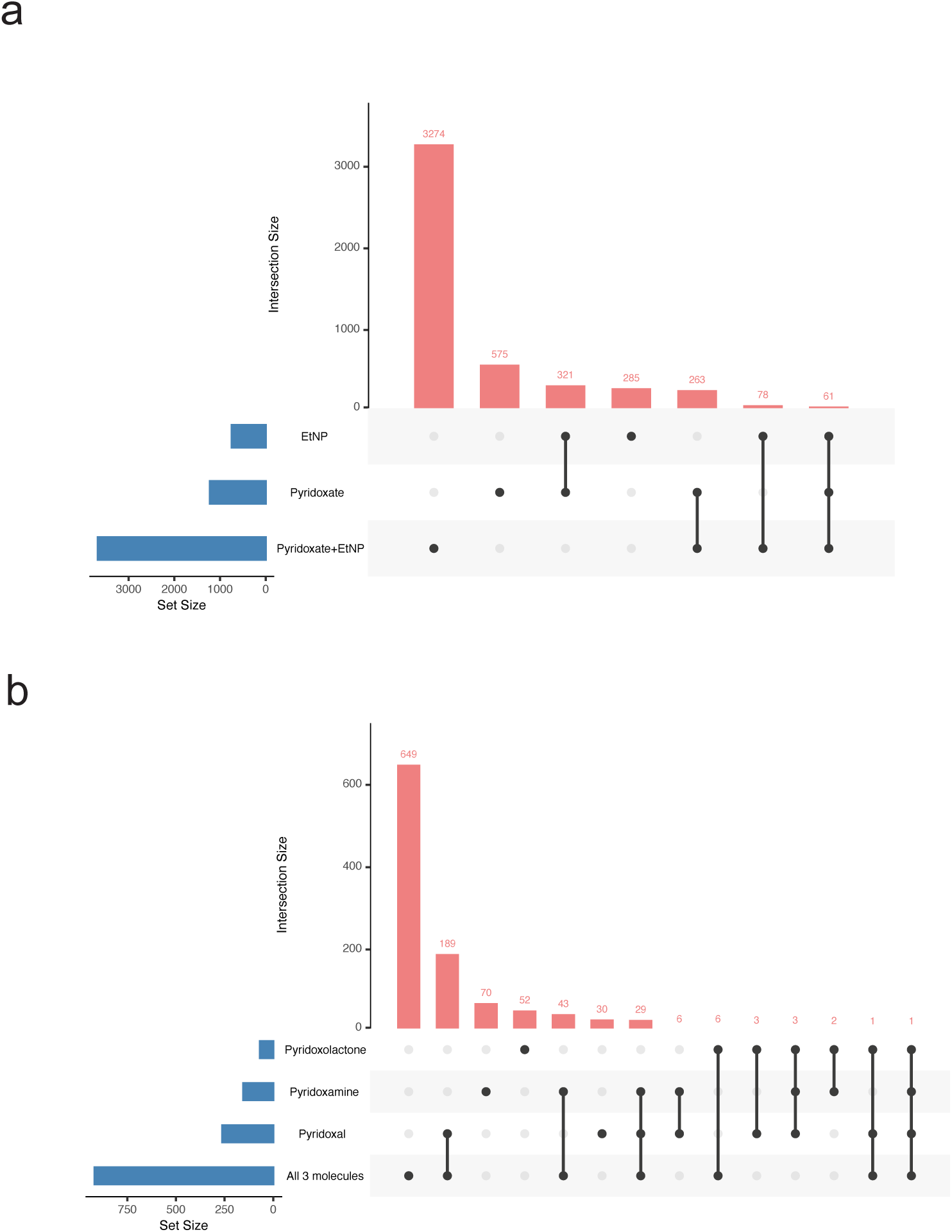
Unique and shared differentially expressed genes for the diatom and bacteria in response to different signals. Upset plots for DEGs in (a) *A. glacialis* exposed to 4-pyridoxate, EtNP, or both, and (b) *P. pseudonitzschiae* exposed to 4-pyridoxolactone, pyridoxamine, pyridoxal, or all three.

**Extended Data Figure 4.**
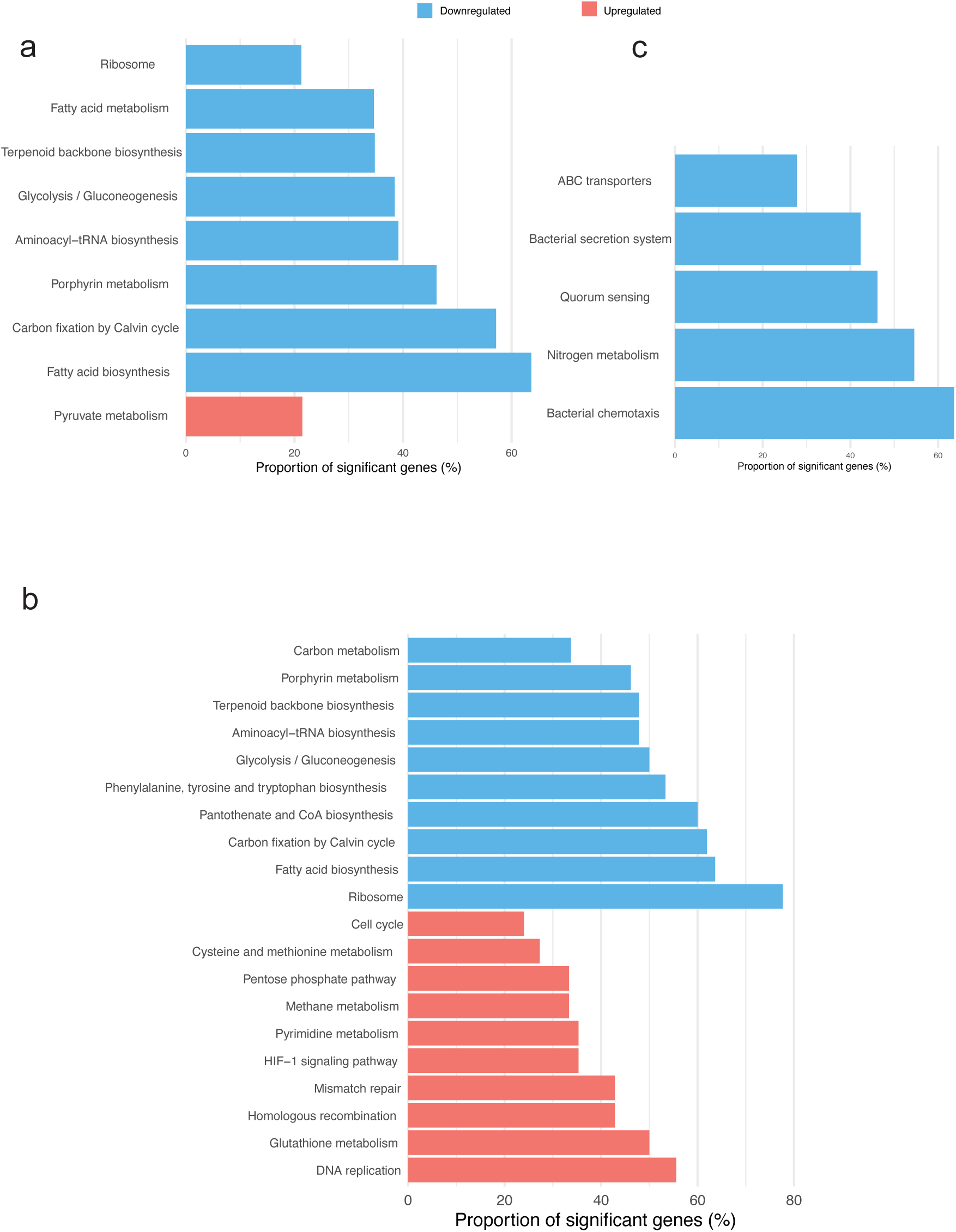
Enriched pathways for signaling between the diatom and the bacterium. Enriched KEGG pathways (p_adj_ < 0.05) of up- and down-regulated genes for *A. glacialis* in the presence of 4-pyridoxate and PEtN at (a) 1 hour, (b) 3 hours, or for (c) *P. pseudonitzschiae* in the presence of 4-pyridoxolactone, pyridoxamine and pyridoxal at 3 hours.

**Extended Data Figure 5.**
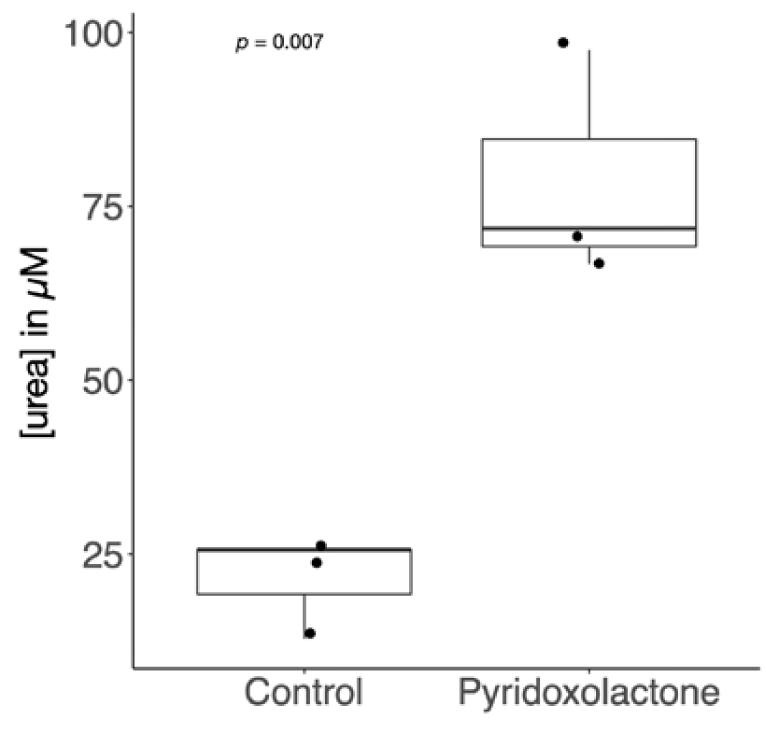
Urea production by *P. pseudonitzschiae* in the presence of 4-pyridoxolactone. Measurements were conducted on biological triplicates.

**Extended Data Figure 6.**
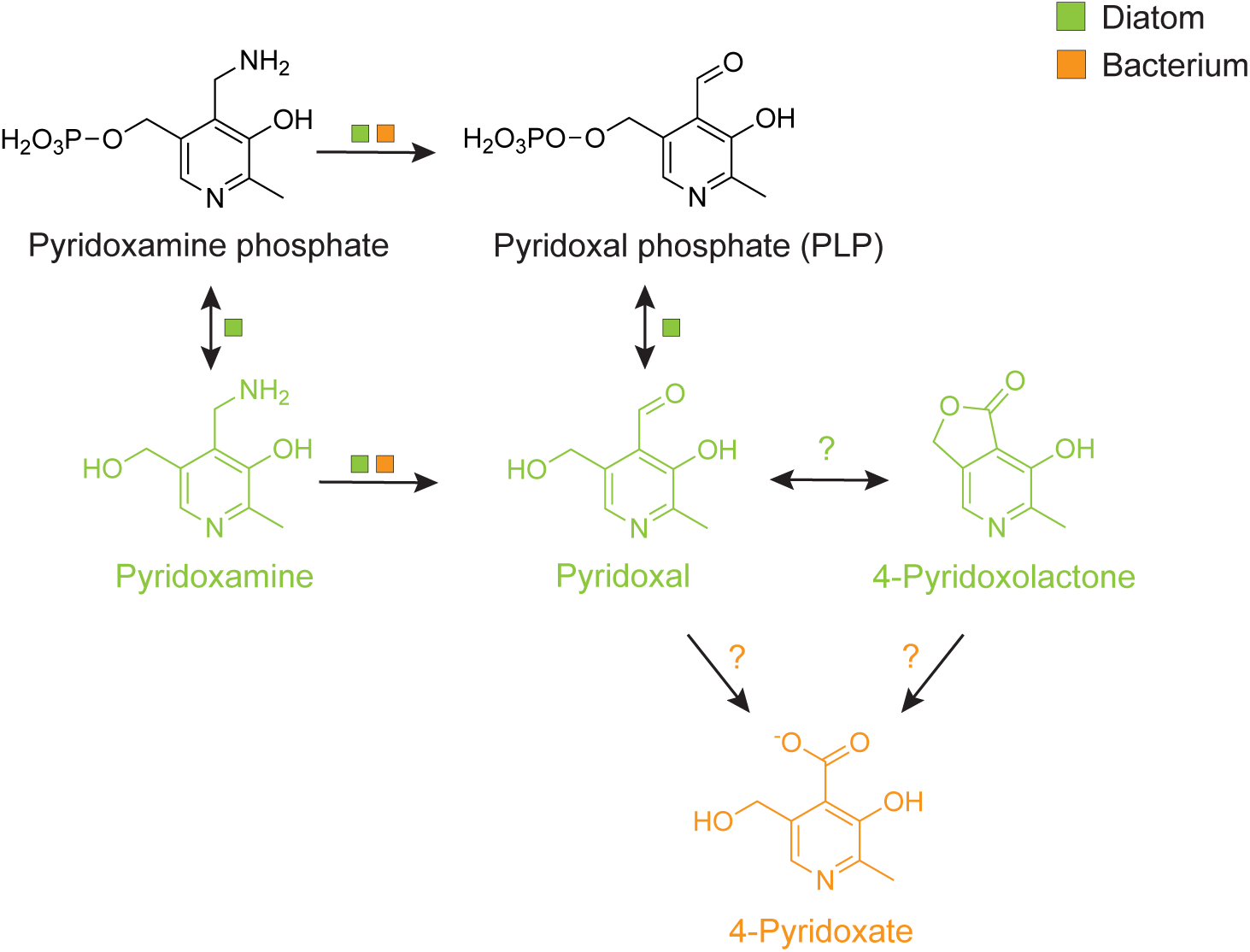
Vitamin B_6_ metabolism in *A. glacialis* and *P. pseudonitzschiae.* Diatom-derived signals are shown in green, while bacterial-derived signals are shown in orange. Boxes indicate presence of genes in either the diatom or bacterial genomes; ‘?’ indicates the lack of a clear homolog in either bacterial or diatom genomes.

**Extended Data Figure 7.**
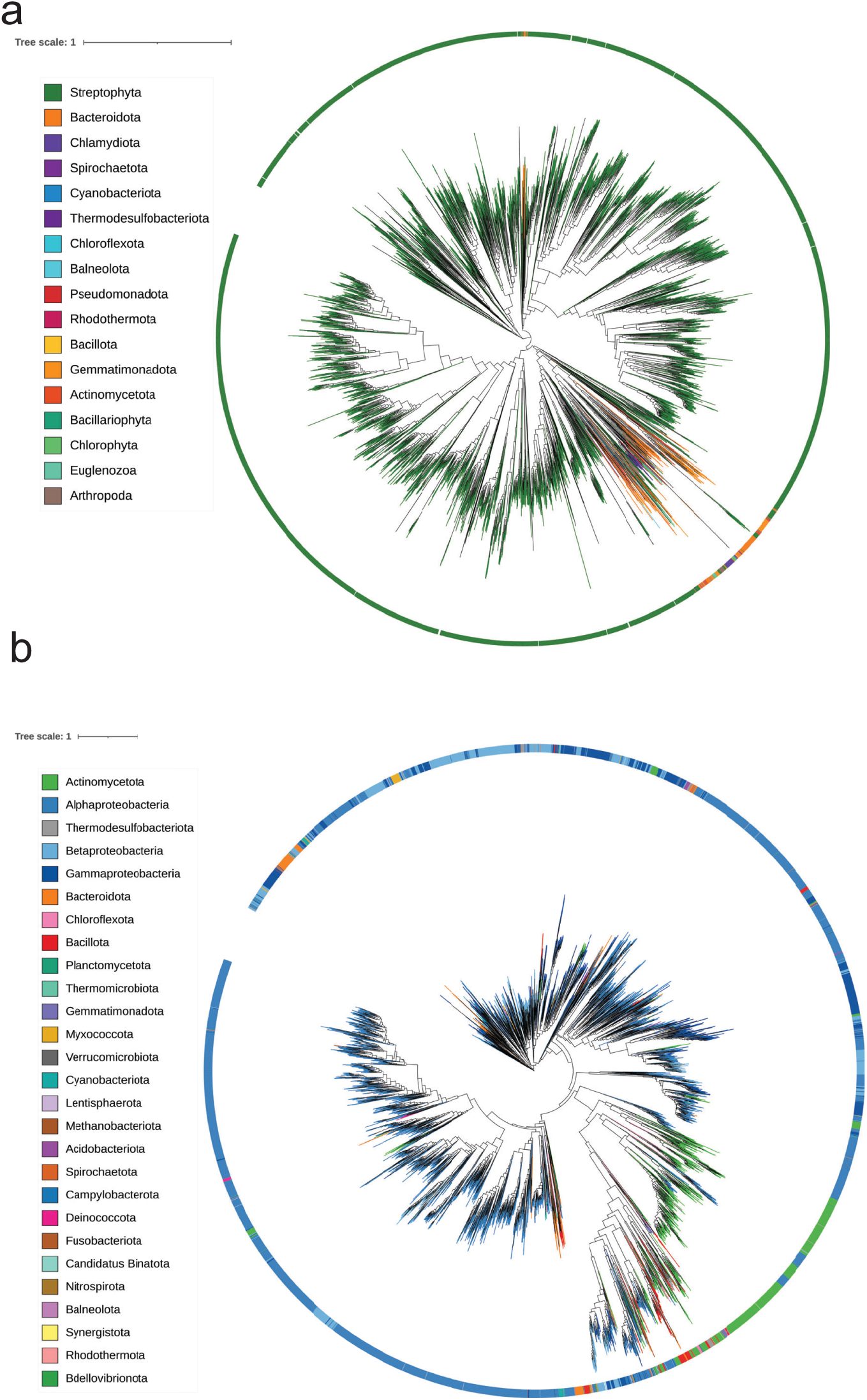
Prevalence of PLDH and PDLA in RefSeq. (a) Distribution of PLDH in RefSeq spanning both eukaryotes and prokaryotes. (b) Distribution of PDLA in RefSeq restricted only to prokaryotes. All IDs and their taxonomy are listed in Supplementary Information Tables 23 and 24.

**Extended Data Figure 8.**
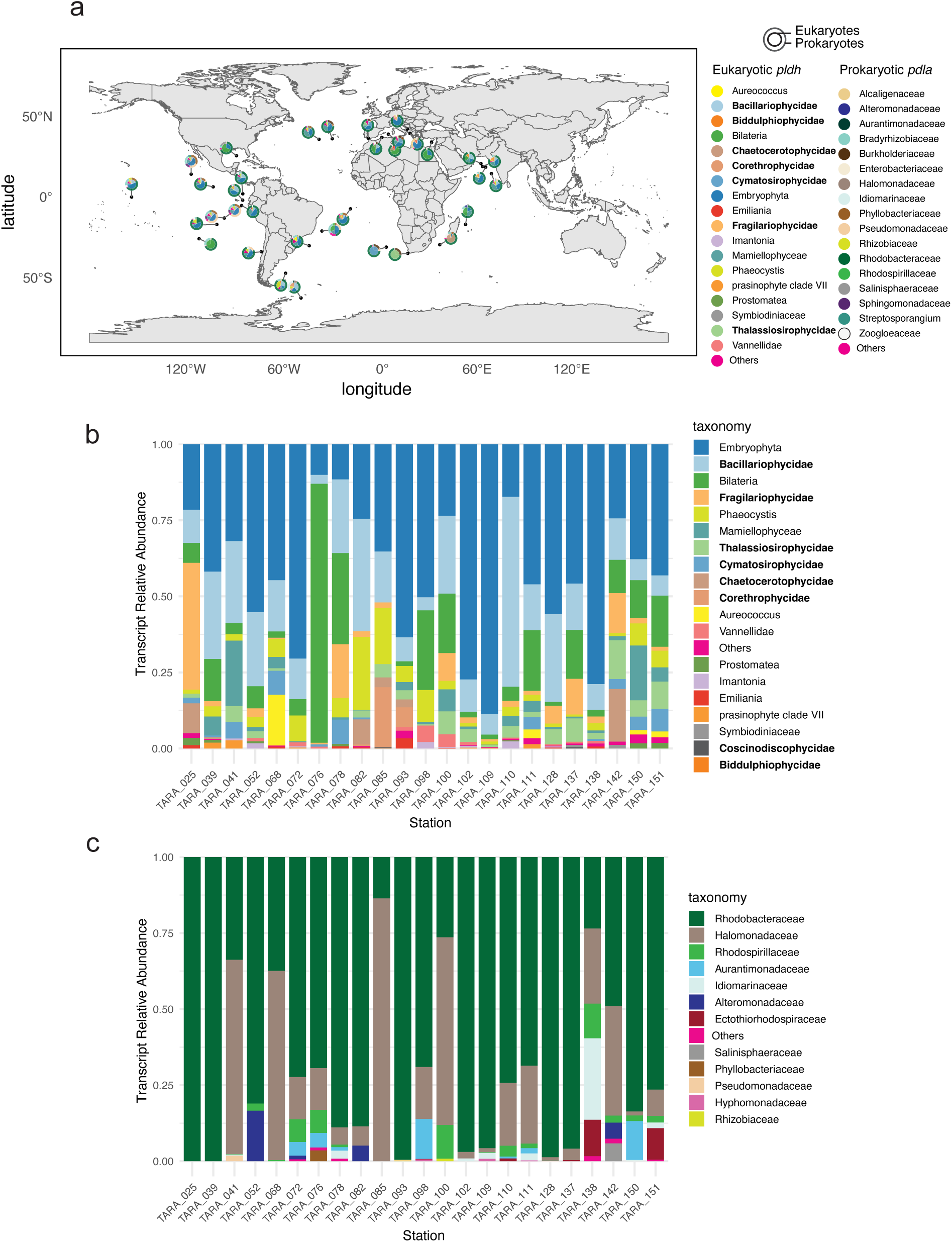
Abundance of *pldh* and *pdla* genes and transcripts in the deep chlorophyll maximum (DCM) across the ocean. HMM profile of each protein sequence was used to mine the Tara Oceans database DCM samples across all available size fractions. (a) Relative abundance and co-localization of *pldh* and *pdla* in Tara metagenomes in the DCM. (b) Transcript relative abundance of *pldh* in Tara metatranscriptomes in the DCM. All bold-face taxa are diatoms. (c) Transcript relative abundance of *pdla* in the same stations as (b) in Tara metatranscriptomes in the DCM.

**Extended Data Table 1.** A summary of the number of differentially expressed genes (DEG) for *Pseudosulfitobacter pseudonitzschiae* and *Asterionellopsis glacialis* in response to compound addition at different time points with *p*-adjusted value <0.05 and log_2_FC >±1 cutoffs. Percent values represent the ratio of genes in each category to the total number of CDS in each genome.

| Organism | Molecule | Incubation time (min) | No. DEG ($p_{\text{adj}} < 0.05$; $\log_2\text{FC} > \pm 1$) | percentage of total CDS |
| --- | --- | --- | --- | --- |
| **Bacterium**<br>(*Pseudosulfitobacter pseudonitzschiae* F5) | Pyridoxine | 60 | 1 | 0 |
|  |  | 180 | 0 | 0 |
|  | Pyridoxamine | 60 | 42 | 0.86 |
|  |  | 180 | 112 | 2.31 |
|  | 4-Pyridoxolactone | 60 | 49 | 1.0 |
|  |  | 180 | 32 | 0.66 |
|  | Pyridoxal | 60 | 257 | 5.3 |
|  |  | 180 | 4 | 0.08 |
|  | Pyridoxamine + 4-Pyridoxolactone + Pyridoxal | 60 | 24 | 0.50 |
|  |  | 180 | 897 | 18.5 |
| **Diatom** (*Asterionellopsis glacialis* A3) | 4-Pyridoxate | 60 | 788 | 4.9 |
|  |  | 180 | 728 | 4.6 |
|  | Phosphoethanolamine | 60 | 710 | 4.4 |
|  |  | 180 | 44 | 0.3 |
|  | 4-Pyridoxate + Phosphoethanolamine | 60 | 1893 | 11.8 |
|  |  | 180 | 3293 | 20.6 |

## References

1. Cho, B. C. & Azam, F. Major role of bacteria in biogeochemical fluxes in the ocean’s interior. Nature 332, 441–443 (1988).

2. Moran, M. A. et al. The Ocean’s labile DOC supply chain. Limnol Oceanogr (2022) doi:10.1002/lno.12053.

3. Moran, M. A. et al. Microbial metabolites in the marine carbon cycle. Nat Microbiol 7, 508–523 (2022).

4. Burg, R. W. & Snell, E. E. The Bacterial Oxidation of Vitamin B6 VI. Pyridoxal Dehydrogenase and 4-Pyridoxolactonase. J. Biol. Chem. 244, 2585–2589 (1969).

5. Seymour, J. R., Amin, S. A., Raina, J.-B. & Stocker, R. Zooming in on the phycosphere: the ecological interface for phytoplankton–bacteria relationships. Nat Microbiol 2, 17065 (2017).

6. Amin, S. A., Parker, M. S. & Armbrust, E. V. Interactions between diatoms and bacteria. Microbiology and Molecular Biology Reviews 76, 667–684 (2012).

7. Bell, W. & Mitchell, R. Chemotactic and growth responses of marine bacteria to algal extracellular products. The Biological Bulletin 143, 265–277 (1972).

8. Focardi, A. et al. Defining the ecological strategies of phytoplankton associated bacteria. Nat. Commun. 16, 6363 (2025).

9. Shibl, A. A. et al. Diatom modulation of select bacteria through use of two unique secondary metabolites. Proc National Acad Sci 117, 27445–27455 (2020).

10. Shibl, A. A. et al. Molecular mechanisms of microbiome modulation by the eukaryotic secondary metabolite azelaic acid. eLife 12, (2024).

11. Fei, C. et al. Quorum sensing regulates ‘swim-or-stick’ lifestyle in the phycosphere. Environ. Microbiol. 22, 4761–4778 (2020).

12. Bronk, D. A. & Glibert, P. M. Application of a 15N tracer method to the study of dissolved organic nitrogen uptake during spring and summer in Chesapeake Bay. Mar. Biol. 115, 501–508 (1993).

13. Bergström, C. A. S., Charman, W. N. & Porter, C. J. H. Computational prediction of formulation strategies for beyond-rule-of-5 compounds. Adv. Drug Deliv. Rev. 101, 6–21 (2016).

14. Mooney, S., Leuendorf, J.-E., Hendrickson, C. & Hellmann, H. Vitamin B6: A Long Known Compound of Surprising Complexity. Molecules 14, 329–351 (2009).

15. Funami, J. et al. 4-Pyridoxolactonase from a symbiotic nitrogen-fixing bacterium Mesorhizobium loti: Cloning, expression, and characterization. Biochim. Biophys. Acta (BBA) - Proteins Proteom. 1753, 234–239 (2005).

16. Mukherjee, T., McCulloch, K. M., Ealick, S. E. & Begley, T. P. Cofactor catabolism II. in Comprehensive Natural Products II (eds. Liu, H.-W. & Mander, L.) 649–674 (Elsevier, 2010). doi:10.1016/b978-008045382-8.00153-2.

17. Pavlovic, Z. & Bakovic, M. Regulation of Phosphatidylethanolamine Homeostasis—The Critical Role of CTP:Phosphoethanolamine Cytidylyltransferase (Pcyt2). Int. J. Mol. Sci. 14, 2529–2550 (2013).

18. Tsimilli-Michael, M. et al. Synergistic and antagonistic effects of arbuscular mycorrhizal fungi and Azospirillum and Rhizobium nitrogen-fixers on the photosynthetic activity of alfalfa, probed by the polyphasic chlorophyll a fluorescence transient O-J-I-P. Appl. Soil Ecol. 15, 169–182 (2000).

19. Gao, Y. et al. Time Series Transcriptome Analysis in Medicago truncatula Shoot and Root Tissue During Early Nodulation. Front. Plant Sci. 13, 861639 (2022).

20. Mohamed, A. R. et al. Dual RNA-sequencing analyses of a coral and its native symbiont during the establishment of symbiosis. Mol. Ecol. 29, 3921–3937 (2020).

21. Kono, M., Kon, Y., Ohmura, Y., Satta, Y. & Terai, Y. In vitro resynthesis of lichenization reveals the genetic background of symbiosis-specific fungal-algal interaction in Usnea hakonensis. BMC Genom. 21, 671 (2020).

22. Via, V. D., Narduzzi, C., Aguilar, O. M., Zanetti, M. E. & Blanco, F. A. Changes in the Common Bean Transcriptome in Response to Secreted and Surface Signal Molecules of Rhizobium etli. Plant Physiol. 169, 1356–1370 (2015).

23. Ge, L. & Seah, S. Y. K. Heterologous Expression, Purification, and Characterization of an l-Ornithine N5-Hydroxylase Involved in Pyoverdine Siderophore Biosynthesis in Pseudomonas aeruginosa. J. Bacteriol. 188, 7205–7210 (2006).

24. Amin, S. A. et al. Photolysis of iron–siderophore chelates promotes bacterial–algal mutualism. Proc National Acad Sci 106, 17071–17076 (2009).

25. Yokochi, N., Nishimura, S., Yoshikane, Y., Ohnishi, K. & Yagi, T. Identification of a new tetrameric pyridoxal 4-dehydrogenase as the second enzyme in the degradation pathway for pyridoxine in a nitrogen-fixing symbiotic bacterium, Mesorhizobium loti. Arch. Biochem. Biophys. 452, 1–8 (2006).

26. Jumper, J. et al. Highly accurate protein structure prediction with AlphaFold. Nature 596, 583–589 (2021).

27. Corbella, M. et al. Catalytic Redundancies and Conformational Plasticity Drives Selectivity and Promiscuity in Quorum Quenching Lactonases. JACS Au 4, 3519–3536 (2024).

28. Kraemer, K., Semba, R. D., Eggersdorfer, M. & Schaumberg, D. A. Introduction: The Diverse and Essential Biological Functions of Vitamins. Ann. Nutr. Metab. 61, 185–191 (2012).

29. Wienhausen, G. et al. Ligand cross-feeding resolves bacterial vitamin B12 auxotrophies. Nature 1–7 (2024) doi:10.1038/s41586-024-07396-y.

30. Banerjee, R. & Ragsdale, S. W. THE MANY FACES OF VITAMIN B12: CATALYSIS BY COBALAMIN-DEPENDENT ENZYMES 1. Annu. Rev. Biochem. 72, 209–247 (2003).

31. Percudani, R. & Peracchi, A. A genomic overview of pyridoxal-phosphate-dependent enzymes. EMBO Rep. 4, 850–854 (2003).

32. Percudani, R. & Peracchi, A. The B6 database: a tool for the description and classification of vitamin B6-dependent enzymatic activities and of the corresponding protein families. BMC Bioinform. 10, 273 (2009).

33. Fitzpatrick, T. B. et al. Two independent routes of de novo vitamin B6 biosynthesis: not that different after all. Biochem. J. 407, 1–13 (2007).

34. Stanulović, M., Jeremić, V., Leskovac, V. & Chaykin, S. New Pathway of Conversion of Pyridoxal to 4-Pyridoxic Acid. Enzyme 21, 357–369 (1976).

35. Kaiser, J. P., Feng, Y. & Bollag, J. M. Microbial metabolism of pyridine, quinoline, acridine, and their derivatives under aerobic and anaerobic conditions. Microbiol. Rev. 60, 483–498 (1996).

36. Salvo, M. L. di, Contestabile, R. & Safo, M. K. Vitamin B6 salvage enzymes: Mechanism, structure and regulation. Biochim. Biophys. Acta (BBA) - Proteins Proteom. 1814, 1597–1608 (2011).

37. Cornforth, D. M. et al. Combinatorial quorum sensing allows bacteria to resolve their social and physical environment. Proc. Natl. Acad. Sci. 111, 4280–4284 (2014).

38. Thomas, S., El-Zayat, A. S., Gurney, J., Rattray, J. & Brown, S. P. Quantitative modeling of multi-signal quorum-sensing maps environment to bacterial regulatory responses. PLOS Biol. 23, e3003316 (2025).

39. Sun, J. et al. Crosstalk between jasmonic acid, ethylene and Nod factor signaling allows integration of diverse inputs for regulation of nodulation. Plant J. 46, 961–970 (2006).

40. Helliwell, K. E., Shibl, A. A. & Amin, S. A. The Molecular Life of Diatoms. in The Molecular Life of Diatoms (eds. Falciatore, A. & Mock, T.) 679–712 (Springer Nature, 2022). doi:10.1007/978-3-030-92499-7_23.

41. Wang, D., Yang, S., Tang, F. & Zhu, H. Symbiosis specificity in the legume-rhizobial mutualism. Cellular microbiology 14, 334–342 (2012).

42. Spaepen, S., Vanderleyden, J. & Remans, R. Indole-3-acetic acid in microbial and microorganism-plant signaling. FEMS Microbiology Reviews 31, 425–448 (2007).

