## Supplementary material for "Vitamin B_6_ Catabolites Act as Interkingdom Signals Within the Marine Microbiome": SI Discussion

**Supplementary Discussion**

**Diatom Metabolites Taken Up by *Pseudosulfitobacter pseudonitzschiae.*** Among the 53 putative diatom metabolites taken up by the bacterium, many were essential amino acids (e.g., Met, Phe, Ile, Tyr, Val, Asp, Lys, His, Tyr) and derivatives, including several functionalized amino acids and derivatives (acetylputrescine, N-amidino-Asp, N-methyl-Asp, 3-methyl-His, N-acetyl-Phe). Surprisingly, the list included many indole and tryptophan derivatives, including IAA, indole-3-acetaldehyde, indole-3-pyruvate, indole-3-carboxylate, tryptamine, and indole, among others. IAA has been shown to be produced by bacteria associated with diatoms and coccolithophores[^1,2^](https://app.readcube.com/library/28bec456-71ba-47f0-b26f-4659a8f9c05b/all?uuid=3526414142385742&item_ids=28bec456-71ba-47f0-b26f-4659a8f9c05b:d951b273-061f-4280-becf-88f961db0c3a,28bec456-71ba-47f0-b26f-4659a8f9c05b:ead80125-d0fc-4d9e-8ea7-259d9591a32b) and a closely related strain to *P. pseudonitzschiae*, SA11, was particularly shown to influence diatom cell cycle through IAA production[^1^](https://app.readcube.com/library/28bec456-71ba-47f0-b26f-4659a8f9c05b/all?uuid=40344475478742015&item_ids=28bec456-71ba-47f0-b26f-4659a8f9c05b:d951b273-061f-4280-becf-88f961db0c3a). Interestingly, hormones with important roles in animals were also observed, though their role in microbes is unclear. These included melatonin, serotonin and lumichrome. Several nucleotides were also observed, such as guanine, adenine and uracil. It is not clear whether these nucleotides were specifically secreted or were a byproduct of sample processing or non-specific cell lysis. Several other metabolites were cryptic in that they do not have a clear role in metabolism, yet they were taken up by the bacterium. These include N-amidino aspartate, N-methyl aspartate, hippuric acid, tryptophanamide, indole-3-carboxylate, hydroxypyridine, 3-methyl-2-oxindole, benzylamine and quinoline.

**Diatom transcriptional response to phosphoethanolamine (PEtN).** After 60 mins of addition of PEtN to *A. glacialis* the diatom upregulated and downregulated 568 and 142 genes, respectively, of which only 139 genes had KO assignments (Supplementary Information Table 16). As typical with diatom genomes, the vast majority of these differentially expressed genes had no annotation and thus interpretation based on these genes is not feasible. For the genes with KO annotations, PEtN addition upregulated molecular systems required for cell division and nucleotide biosynthesis. Specifically, purine biosynthesis was activated as evidenced by the upregulation of several enzymes in the purine metabolism pathway, including functions involved in increasing guanine nucleotide supply (GMP synthase, K01951), correcting mutagenic or non-canonical nucleotides incorporation into DNA synthesis during metabolic shifts (XTP/dITP diphosphohydrolase, K01519), and global transcriptional reprogramming via cAMP (adenylate cyclase, K01768) that can switch on nitrogen assimilation and carbon fixation and cell cycle genes[^3^](https://app.readcube.com/library/28bec456-71ba-47f0-b26f-4659a8f9c05b/all?uuid=5650820269839016&item_ids=28bec456-71ba-47f0-b26f-4659a8f9c05b:638c1e4b-a62e-4c51-9e61-df4a1447005a). These patterns are coupled with upregulation of several cell cycle genes (K03456, K04683, and K09392) that prepare the cell for division and also to several spliceosome genes (K12815, K12831, K12852) that likely process transcriptome shift due to PEtN. In addition within purine metabolism, the diatom also activates allantoicase (K01477) that may breakdown nitrogen indirectly from PEtN-nitrogen converted to purines to generate urea, adding to the urea pool that is supplied by the bacteria. These patterns are coupled with several upregulation patterns in amino sugar and nucleotide sugar metabolism, where galactose is converted to UDP-galactose via galactokinase (K00849) and UDP-sugar pyrophosphorylase (K12447) to boost the UDP-sugars pool in the cell for glycolipid and polysaccharide biosynthesis. These patterns broadly point to an increase in nucleic acid biosynthesis, potentially to prepare the cell for division, and membrane remodeling. The latter is reinforced by upregulation of sulfate conversion to APS and further to PAPS (K13811), which may provide sulfated lipids or polysaccharides to the cell in the presence of PEtN, while the former is reinforced by upregulation of several chromatin remodeling genes that are linked to cell proliferation and DNA synthesis (K11251, K11320, K05610, and K11661).

Interestingly, select canonical photosynthesis genes constituting Photosystem I and II were downregulated (*psaA* K02689, *psaB* K02690, *psbA* K02703 and *psbC* K02705) along with Rubisco (K01602) due to one of several possibilities: activation of DNA replication, cell cycle genes and chromatin remodeling may signal that PEtN is interfering with light-regulated cell synchrony; remodeling of membranes, especially the thylakoid membrane may interfere with photosynthesis and trigger this pattern. These patterns partially overlap with patterns of downregulation of carbon fixation and autotrophy when the diatom is supplemented with PEtN and 4-pyridoxate.

In general, diatom-derived metabolites metabolism were not strongly affected by PEtN except for methionine, which the bacterium took up. Methionine biosynthesis gene (K00548) that converts homocysteine to methionine exhibited one of the highest Log_2_FC values in the transcriptome at 60 mins, log_2_FC = 6.0 (Supplementary Information Table 16). At 180 mins post PEtN addition, differential expression was majorly muted with only 14 and 30 genes up- and downregulated with the vast majority lacking annotations (Supplementary Information Table 17).

**Diatom transcriptional response to 4-pyridoxate (4-PA).** After 60 mins of addition of 4-PA to *A. glacialis* the diatom upregulated and downregulated 643 and 145 genes, respectively, of which only 121 genes had KO assignments (Supplementary Information Table 14). Similar to the response in the presence of PEtN, several reactions and pathways were regulated. For example, purine metabolism was differentially regulated with adenylate cyclase (K01768) being upregulated again, conversion of sulfate to PAPS and activation of allantoicase, which presumably can generate urea from N-rich molecules like purines. Likewise, amino sugar and nucleotide sugar metabolism genes were generally upregulated with 4-PA as they were with PEtN, with galactose conversion to UDP-galactose. Exposure to 4-PA further activated more genes in this pathway to generate UDP-glucose, UDP-glucuronate, UDP-xylose, UDP-arabinose and UDP-N-acetyl-galactosamine (K00849, K12447, K18674). These amino-sugars may be involved in biosynthesizing sulfated galactans, particularly as a sulfotransferase genes are highly upregulated in the transcriptome (K01016, K01025), which may play a role in diatom cell membranes. Cell cycle genes and spliceosome genes were also upregulated as seen with PEtN, indicating an overlap of regulation of both these signals.

Diatom-derived metabolites metabolism was generally more evident with 4-PA than in PEtN with branched amino acid degradation generally downregulated (K00248, K07511, K07508), concomitant with secretion of these metabolites by the diatom for the bacterium. An earlier step in the pathway is upregulated (K09699), suggesting potential accumulation of branched-amino acids-coA intermediates. Some of the downregulated enzymes in this pathway are also involved in β-oxidation of fatty acids (K00248, K07511, K07508, in addition to K07515), which is also downregulated along with fatty acid elongation (K07508, K07511, K07515 and K10258). These patterns suggest a concerted curbing of ROS overproduction. Several genes in the carbon fixation pathway were upregulated, particularly glyceraldehyde 3-phosphate dehydrogenase (K00134), which provides ATP/NADH^+^ to compensate for the downregulation of β-oxidation.[^4^](https://app.readcube.com/library/28bec456-71ba-47f0-b26f-4659a8f9c05b/all?uuid=6540979231705157&item_ids=28bec456-71ba-47f0-b26f-4659a8f9c05b:8e524103-2826-422e-a845-81deabcc3a13) In contrast to PEtN, 4-PA upregulates photosynthesis genes, such as *petD* (K02637), *psbO* (K02716), *psbU* (K02719), and *psbQ* (K08901). Curiously, the PEtN and 4-PA transcriptome shows no differential expression of these genes (Supplementary Information Tables 18-19).

At 180 mins, in general these patterns persisted in some of the aforementioned pathways with minor differences in the genes that are differentially expressed or pathways that appear to show minor mixed regulation (i.e., from purely upregulated to mostly upregulated). Some pathways showed significantly dampened regulation (e.g., branched-amino acid degradation) and others were not differentially regulated, particularly β-oxidation of fatty acids and fatty acid elongation. The main exceptions to these patterns were photosynthesis and carbon fixation, which showed downregulation at this time point. For example, Rubisco showed downregulation, despite not being differentially expressed at 60 mins. Several photosynthesis genes were downregulated though these genes were not differentially expressed at 60 mins, such as *petF* (K02639), *psaA* (K02689), *psaB* (K02690), *psbA* (K02703) and *psbB* (K02705). These patterns are consistent with the persistent levels of cAMP, inferred from the continual upregulation of adenylate cyclase, which have been shown to repress photosynthesis[^5–7^](https://app.readcube.com/library/28bec456-71ba-47f0-b26f-4659a8f9c05b/all?uuid=284474343320149&item_ids=28bec456-71ba-47f0-b26f-4659a8f9c05b:3f5423f0-97fd-4be4-ae08-6f4035741303,28bec456-71ba-47f0-b26f-4659a8f9c05b:83dd0fe6-4e40-4bb9-9131-c969a7a675bc,28bec456-71ba-47f0-b26f-4659a8f9c05b:26bad1e3-187c-4c34-b1ba-38685ff97c60).

**Bacterial transcriptional response to pyridoxamine.** After 60 mins, *P. pseudonitzschiae* upregulated genes involved in the biosynthesis of the amino acids valine and isoleucine (K01653) and the catabolism of glutamine (K13821). After 180 mins, the bacterium upregulated branched amino acid ABC transporters (K01995, K01997, K01998) though amino acid biosynthesis/catabolism genes were generally muted. Instead, the bacterium upregulated type IV secretion system (K03195, K03199, K03200, K03201, K03204) that make up part of the *tad* gene cluster required for the assembly and secretion of adhesive fimbrial proteins, known as Flp pili. These have been shown to be critical for bacterial attachment to biotic and abiotic surfaces and are highly enriched in phycosphere colonizer bacteria[^8,9^](https://app.readcube.com/library/28bec456-71ba-47f0-b26f-4659a8f9c05b/all?uuid=7834808549592956&item_ids=28bec456-71ba-47f0-b26f-4659a8f9c05b:9e206206-4f5c-4acc-8d35-bfaf8d62591d,28bec456-71ba-47f0-b26f-4659a8f9c05b:4b28fa0d-221c-4039-9487-dda2b997c247). The breakdown of creatine, a metabolite taken up by the bacterium from *A. glacialis*, to urea was also upregulated (K08688), coinciding with urea excretion as discussed in the main text (Fig. 3, Extended Data Fig. 5).

**Bacterial transcriptional response to pyridoxal.** Most DEGs were found at 60 mins, with the 180 mins transcriptome showing only 4 DEGs. At 60 mins, the bacterium mainly upregulated catabolism of diatom metabolites it anticipates to take up, including valine, leucine (K00249), 5-methylcytosine (K01485) and hypoxanthine (K11178). It also upregulated genes for the conversion between creatine and creatinine (K01470, K01485, K01473, K01474), both of which were taken up from the diatom. The breakdown of creatine generates urea, and this gene was upregulated with other signals; however, it was not differentially expressed with pyridoxal. Related to the urea cycle, the bacterium also upregulated the conversion of N-acetylornithine to ornithine from the diatom and N-acetylcitrulline to citrulline (K01438). Several genes related to chemotaxis were upregulated (K00575, K03406, K03407, K03408, K03413), potentially indicating that with just pyridoxal the diatom may be interfering with phycosphere colonization of potential cheaters who may be sensing pyridoxal alone. Type IV secretion systems were not as strongly induced as with pyridoxamine, with only two genes upregulated (K03201, K03205). In strong contrast to downregulation of nitrate assimilation and reduction with the 3 signals, pyridoxal alone upregulated nitrate transport and reduction to nitrite (K00372, K15576), presumably distinguishing the response of cheaters perceiving only pyridoxal who would still require inorganic nitrogen to grow from symbionts who would rely on DON from the diatom.

**Bacterial transcriptional response to 4-pyridoxolactone.** With only dozens of differentially expressed genes, 4-pyridoxolactone regulation of the bacterial response is mostly muted. One exception at 60 mins is the production of urea via the conversion of arginine to ornithine (K01476) is upregulated and coincides with the bacterial enhanced production of urea in the presence of pyridoxolactone (Extended Data Fig. 5).

**References**

[1. Amin, S. A. *et al.* Interaction and signalling between a cosmopolitan phytoplankton and associated bacteria. *Nature* **522**, 98–101 (2015).
2. Segev, E. et al. Dynamic metabolic exchange governs a marine algal-bacterial interaction. *eLife* **5**, 514 (2016).
3. Watzer, B. et al. The Signal Transduction Protein PII Controls Ammonium, Nitrate and Urea Uptake in Cyanobacteria. *Front. Microbiol.* **10**, 1428 (2019).
4. Wang, S. et al. The Role of Glyceraldehyde-3-Phosphate Dehydrogenases in NADPH Supply in the Oleaginous Filamentous Fungus Mortierella alpina. Front. Microbiol. **11**, 818 (2020).
5. Mantovani, O., Haffner, M., Selim, K. A., Hagemann, M. & Forchhammer, K. Roles of second messengers in the regulation of cyanobacterial physiology: the carbon-concentrating mechanism and beyond. *microLife* **4**, uqad008 (2023).
6. Tanaka, A., Ohno, N., Nakajima, K. & Matsuda, Y. Light and CO2/cAMP Signal Cross Talk on the Promoter Elements of Chloroplastic β-Carbonic Anhydrase Genes in the Marine Diatom Phaeodactylum tricornutum. *Plant Physiol.* **170**, 1105–1116 (2015).
7. Ohmori, M. & Okamoto, S. Photoresponsive cAMP signal transduction in cyanobacteria. *Photochem. Photobiol. Sci.* **3**, 503–511 (2004).
8. Bentzmann, S. de, Aurouze, M., Ball, G. & Filloux, A. FppA, a Novel Pseudomonas aeruginosa Prepilin Peptidase Involved in Assembly of Type IVb Pili. *J. Bacteriol.* **188**, 4851–4860 (2006).
9. Isaac, A., Francis, B., Amann, R. I. & Amin, S. A. Tight Adherence (Tad) Pilus Genes Indicate Putative Niche Differentiation in Phytoplankton Bloom Associated Rhodobacterales. Front. Microbiol. **12**, 718297 (2021).](https://app.readcube.com/library/?style=Nature+%7B%22language%22:%22en-US%22%7D)
