## Supplementary material for "Vitamin B_6_ Catabolites Act as Interkingdom Signals Within the Marine Microbiome": SI file

**Supplementary Methods**

***Chemicals.*** All chemicals (Optima-Grade or similar) were purchased from ThermoFisher Scientific (Waltham, US). Formic acid, Na^15^NO_3_ and all vitamin B_6_ related metabolites were purchased from MERCK (Darmstadt, Germany), and reference compounds from the IROA-LSMS and IROA-IS kits were purchased from IROA technologies (Nellysford, US).

***Diatom and bacterial growth.*** Axenic *A. glacialis* strain A3 culture (CCMP3542) was maintained in *f/2*+Si medium^1^ with Na^15^NO_3_ as the nitrogen source in semi-continuous batch cultures^2^. All cultures were incubated in growth chambers (Percival, Perry, US) at 22°C, 130 µE m^-2^ s^-1^ on a 12:12 light/dark cycle. Light flux was measured using a QSL-2100 PAR Sensor (Biospherical Instruments Inc., San Diego, USA). Growth was monitored by measuring *in vivo* fluorescence using a 10-AU fluorometer (Turner Designs, San Jose, CA). The labeled cultures were transferred at least four times using semi-continuous batch cultures over 4-6 weeks for stable labeling.

All bacteria, including *P. pseudonitzschiae* F5, were isolated as described previously^3^ and were grown from -80°C glycerol stocks on ZoBell marine agar plates^4^. Plates were incubated at 25°C until colonies were visible. Bacterial growth was quantified using optical density at 600 nm in ZoBell marine broth.

***Co-culture generation and growth.*** For co-cultures of *A. glacialis* with *Alteromonas macleodii* F12, *Phycobacter azelaicus* F10 and *Pseudosulfitobacter pseudonitzschiae* F5 and for co-cultures of *A. glacialis* with *P. pseudonitzschiae*, bacteria were grown in ZoBell marine broth^4^ in the dark overnight from a single colony at 26°C and shaking at 180 rpm. Overnight cultures were used to inoculate 50 mL of broth that was grown to an optical density (OD_600nm_) of 0.5. Cultures were centrifuged at 4000 rpm for 10 min followed by washing and resuspending twice with sterile f/2+Si medium. Each washed stock was used to inoculate ^15^N-labeled axenic A. glacialis cultures at a density of 5000 cells mL^−1^ and at a total bacterial density of ~1× 10^6^ cells mL^−1^.

***Preparation for NanoSIMS Card-FISH.*** Axenic *A. glacialis* incubated with *P. pseudonitzschiae* or all three bacterial combinations were sampled at 0, 2, 6, and 12 hours post inoculation with bacteria and were quenched with 20% paraformaldehyde solution added directly to the culture vessels to a final concentration of 1‐2 % PFA at room temperature, mixed gently and pH checked to be around pH 7.5. Samples were incubated in paraformaldehyde for 12-18 hours at 4°C after which samples were filtered with a gentle vacuum (<200 mbar) onto Au-Pd sputtered 0.2 µm polycarbonate membrane filters supported by a cellulose nitrate filter. Filters were washed twice with 10 mL sterile MilliQ water, dried by tapping onto blotting paper and left to dry. Once completely dry, filters were stored at ‐20°C until Nano-secondary ion mass spectrometry (NanoSIMS) measurement.

***Microscopic identification, subsample preparations and markings for SIMS.*** The Au-Pd filters were initially scanned for diatom cells using a Laser Micro Dissection (LMD) microscope 6500 (Leica Wetzlar, Germany) fitted with blue (440-85 nm) and green (517-41 nm) excitation filter sets. Briefly, the filter was placed in a 2 mL Eppendorf tube with cells facing inwards and amended with 1 mL sterile 0.1 M PBS buffer. Using a 1 mL pipette, the filter was gently washed multiple times with PBS. Using small aliquots (100–200 μL), the re-suspended cells were gravity filtered onto a new pre-sputtered Au-Pd (0.3 μm pore size, 25mm diameter) and densities were checked. Laser markings and arrows were directly added to the filter to easily re-identify areas for analysis during the SIMS. Additional microscopic images were taken for orientation purposes during the SIMS analyses and for post-processing using look@nanoSIMS^5^.

***CARD-FISH staining and acquisition.*** CARD-FISH was done on similarly prepared sample filters as used for NanoSIMS and labeling with horseradish peroxidase (HRP) labeled probes as described previously in Kitzinger *et al*^6^. In short, cells on Au-Pd filter wedges were treated for endogenous peroxidases, permeabilized with proteases, labeled with Horseradish peroxidase (HRP) probes and HRP signal enhanced by deposition of fluorescent tryptamide. Alphaproteobacteria probes were used to label the Rhodobacteraceae, while *A. macleodii* (Gammaproteobacteria) were DNA stained (below). All samples were washed 2 times with deionized water and dried in a petri dish. To stain diatom/bacteria DNA, filter wedges were incubated in 1 mL Milli-Q water containing 1 µl of 1 µg/µl 4′,6-diamidino-2-phenylindole (DAPI) for 10 min at 4°C. Slides were further washed in deionized water for 5 min and air dried. For ﬂuorescence microscope imaging, images were acquired with an Axio Imager M2 (Zeiss, Oberkochen, Germany) with ZEN-2 (blue version) software using 40× and 100× objectives. The following channels were used to view the images: DAPI for diatom and bacteria (*A. macleodii*) DNA, FITC (EGFP-HC ﬁlter set, Rhodobacteraceae), and bright ﬁeld light.

***SIMS analysis.*** SIMS analyses were performed using a CAMECA NanoSIMS 50 L instrument (Cameca, Gennevilliers, France) or a CAMECA ims1280 large-geometry (LG-) SIMS instrument. Brieﬂy, for analyses performed on the NanoSIMS 50 L, the cells were rastered with a primary ion beam with a current between 1–2 pA and a beam size <100 nm. Mass resolving power (MRP) in all measurements was > 7000. The primary ion beam was used to raster the area of interest at a size between 40 × 40 µm and 50 × 50 µm with an image size of 256 × 256 pixels or 512 × 512 pixels over the chosen raster size with a dwelling time of 1 or 2 ms per pixel. Negative charged secondary ions (^12^C^−^, ^13^C^−^, ^12^C^14^N^−^, ^12^C^15^N^−^, ^31^P^−^, ^32^S^−^) were collected simultaneously in 6 parallel electron multiplier detectors of the multi-collection system of the instrument. All scans (15–200 planes per diatom cell) were corrected for drift of beam and sample stage and accumulated after acquisition using the look@nanoSIMS software package^5^.

Isotope ratio images were created as the ratio of the sum of counts for each pixel over all recorded planes of the investigated isotope and the main isotope. Regions of interest (ROIs) were deﬁned using the secondary electron (SE), ^12^C^14^N^−^, ^31^P^−^ and epi-ﬂuorescent images taken before the analysis. For each ROI, the isotopic ratio was calculated using look@nanoSIMS software package^5^.

The calculations of N-exchange rates for the individual microbial cells was computed using measured ^12^C^15^N^−^ ion signal, as a proxy for ^15^N atoms, with a method adapted from Foster *et al.*^7^ To estimate the uptake ratio the averaged ^12^C^15^N^−^/(^12^C^14^N^−^+^12^C^15^N^−^) ratios at timepoint 2h were divided by timepoint 0h resulting in the labeling factor (LF). To estimate the uptake ratio in mol h^-1^ cell^-1^ the mean ^12^C^15^N^−^ ion numbers of the measured ROIs (cells) at 2h were averaged and normalized with LF and divided by the number of hours.[^6^](https://app.readcube.com/library/28bec456-71ba-47f0-b26f-4659a8f9c05b/all?uuid=8319437388699303&item_ids=28bec456-71ba-47f0-b26f-4659a8f9c05b:72156b82-10b3-4dc5-9333-84ae31b4f24b)

***Harvesting of Diatom spent media and inoculation with P. pseudonitzschiae*.** *A*xenic *A. glacialis* spent media were harvested by filtration from larger cultures. For this 6 x 1 L sterile culturing flasks were prepared with ^15^N-nitrate *f/2*+Si medium and inoculated with 25 mL of ^15^N labelled *A. glacialis* cultures at an average cell density of ~1 × 10^3^ cells mL^−1^. Cultures were grown until mid-exponential growth and then were carefully filtered using 3-µm polycarbonate filters (Whatman, NJ, USA) to remove *A. glacialis* cells. 300 mL of exudate solution was extracted using SPE for bulk analysis to detect and putatively annotate diatom ^15^N labeled metabolites (See below). The rest of the exudate was separated into sterile test tubes (25 mL each) to examine uptake of exudates by the bacterium. The exchange experiment was set up in 2 steps as shown in Figure 2. After preparation of the tubes five test tubes were directly extracted at timepoint 0 hour and used to measure the starting amounts of ^15^N-labeled metabolites, while another set of tubes were incubated for 6 hours to account for abiotic metabolite degradation. Ten test tubes were inoculated with *P. pseudonitzschiae* to a final density of ~1 × 10^6^ cells mL^−1^ and incubated at 22°C in the diatom growth chamber during the light period. After 3 hours all inoculated samples were sterile filtered through 0.2-µm polycarbonate filters (Whatman, NJ, USA) and transferred aseptically into fresh sterile glass test tubes. Five inoculated samples were directly extracted for metabolites using SPE as described below. For Step 2, five filtered exudate samples from Step 1 were inoculated with *A. glacialis* at a cell density of ~7 × 10^4^ cells mL^−1^. After an additional 3 hours, all samples were filtered and metabolites were extracted as described below.

***Recovery of Diatom exudate metabolites.*** To extract (labeled) exudate molecules, the spent media were filtered at each step as described above. Agilent PPL Bond Elute solid-phase extraction (SPE) cartridges (1 mL cc, 200 mg for 30 mL samples, 6 mL cc, 1 g for 300 mL samples) (Agilent Technologies, Santa Clara, USA) were used to extract and concentrate metabolites. The columns were activated according to the manufacturer’s instructions and subsequently used to remove and desalt organic molecules from the ﬁltrates as previously described^8^. Extracts were eluted with 1 mL and 5 mL, respectively, methanol in 0.1% (v/v) formic acid and dried using an evaporator (SpeedVac SC210A; Thermo Savant, Holbrook, NY, USA). Samples were immediately stored at −20°C until FT-ICR-MS analysis.

***Preparation for mass spectrometry.*** The bulk extract, from which diatom metabolites were annotated, was redissolved in 400 µL of 90% MeOH and spiked with 15 µL of a mixture of 343 nitrogen-containing molecules from the MSMLS library (IROA Technologies, Nellysford, USA) (Supplementary Information Table 1), and 15 µL of ^13^C labeled internal standards from the IROA-IS library (IROA Technologies, Nellysford, US), which were used as a reference for the identification of ^15^N labeled metabolites from diatom exudates. The IROA-IS was dissolved in 1.2 mL of MilliQ water according to the manufacturer’s recommendations. Subsequent samples from the above-described exchange experiment were spiked with 5 µL of ^13^C labeled internal standards IROA-IS for calibration.

***FT-ICR-MS metabolic profiling.*** Fourier-transform ion cyclotron resonance mass spectrometry (FT-ICR-MS or FT-MS thereof) allows the measurement of high-resolution accurate mass and isotopic fine structure for direct determination of the chemical formula of a compound^9^. FT-ICR-MS direct infusion signals in electrospray ionization (ESI) source are prone to ion-suppression due to high salt content in the sample matrix; this requires a desalting step using a solid-phase extraction (SPE) step or a similar approach^10^. Therefore, PPL-bond elute SPE (Agilent Technologies, USA) were used as described above. Direct infusion of samples at 1 µL/min were used to acquire high-resolution mass spectra on a Bruker solariX FT-MS (Bruker Daltonics GmbH, GER) equipped with a 7T superconducting magnet (Magnex Scientific Inc., UK) operated in negative ESI mode (ESI-). The mass spectra acquired in ESI- produces a smaller number of adducts, as well as higher peak resolution compared to positive ESI in the FT-ICR-MS. Spectra were acquired with a time domain of 4 mega words over a mass range of *m/z* 100 to 1000, with an optimal mass range from *m/z* 200-600. Three hundred scans were accumulated for each sample. The following MS parameters were used: ionization mode: ESI-, mass range: *m/z* 100 to 1000, capillary voltage: 4500 V, End Plate Offset –700 V, Nebulizer pressure: 2 bar, Dry gas flow (10 L/min), Dry temperature: 220°C. Iontrap: Source Optics, Capillary Exit: –200V, Deflector Plate: –220V, Funnel1: –150V, Skimmer: –15V, Funnel RF App: 150Vpp; Octopole: Frequency: 5 MHz, RF Amplitude: 350 Vpp, Collision Cell: Collision Voltage: –10V, DCextract Bias: –0.6V, RF Frequency: 2 MHz, Collision RF Amplitude: 1500 Vpp, Transfer Optics: TOF: 0.5 ms, Frequency: 4 MHz, RF Amplitude 350 Vpp. Analyzer: ParaCell: Transfer Exit Lens: 23.0 V, Analyzer Entrance: 10.0 V, Side kick: 0.0 V, Side kick Offset: 1.5 V, Front trap plate: –2.0 V, Back Trap plate: –2.0 V, Back trap Quench: 30 V, Sweep excitation: 21%; Shimming DC Bias: 0°: –1.468, 180°: –1.532, 90°: –1.420, 270°:–1.584, ICR Fills: 1. MS/MS parameters collision voltage: 10V, 30V, 50V.

***Data calibration and feature picking.*** The FT-MS mass spectra were internally calibrated in Dataanalysis 5.0 (Bruker Daltonics GmbH, Germany) on primary metabolites (Amino-, fatty-, organic-acids). Peak alignment was performed with maximum error thresholds of 0.01 ppm with a cut-off signal-to-noise ratio (S/N) of 4. The resulting peak tables were exported to peak lists.

***Data annotation.*** To make use of ultrahigh resolution data (250k-1M resolution) of various stable isotope labeled metabolites in complex mixtures, a command line python (3.7) script, *Molecular Isotope Mass Identifier (MIMI) version 1.0 (https://github.com/NYUAD-Core-Bioinformatics/MIMI)*, was employed for calculation of chemical formulas from exact masses^11^. Processing parameters were set to negative ionization mode (-H), a 0.5 ppm threshold or exact mass match (-p). As a reference library, the supplied KEGG chemical database was used.

The bulk *A. glacialis* exudate bulk sample, spiked with MSMLS as described above, was measured in MS and MS/MS mode at 10V, 30V, and 50V collision energies. Chemical formulas were calculated according to natural isotopic abundance and ^15^N in each MS mode. Isotopic fine structure peaks calculated for each label type were used as confirmation of chemical formulas of precursors and MS/MS fragments. Annotations according to MS/MS fragmentation were assisted by the prediction of fragmentation patterns using CFM-ID3.0^12^. In short, the parent compounds were submitted as SMILES^13^ string and prediction was submitted with spectra type ESI, for ESI- and [M-H]^-^ as primary ion type. To verify chemical structures, ^15^N-labeled metabolite candidates were compared with the observed fragmentation patterns of non-labelled internal standards from MSMLS matching only nitrogen-containing fragments at a match ratio ≥80% (Supplementary Information Table 1-3, Extended Data Fig. 2). Abundance estimates for the starting condition were based on direct peak comparison between spiked non-labeled N-containing metabolites and ^15^N-labeled metabolites from *A. glacialis* exudate normalized to the ^13^C labelled metabolites from IROA-IS. Subsequent samples from the exchange experiment were measured only in MS1 at 500,000 – 1M at 110 m/z resolution and chemical formulas were calculated from accurate mass and isotopic fine-structure as described above.

**Diatom incubation experiments.** To test the transcriptional response of *A. glacialis* to EtNP, 4-pyridoxate (4-PA) or a mixture of both metabolites (Merck, Darmstadt, Germany), cells were grown to a starting density of 8 x 10^5^ cells mL^-1^ in sterile f/2 media. Cultures were supplemented with a final concentration 50 μM filter-sterilized solution of each metabolite individually or 25 μM of both metabolites when combined. Controls were supplied with an equal volume of solvent necessary to dissolve the metabolite of interest. All samples were done in triplicates.

***Diatom RNA isolation and sequencing.*** For eukaryotic RNA, cells were filtered through 3-µm polycarbonate filters (43 mm, Whatman, NJ, USA). All filters were flash-frozen in liquid nitrogen and later stored in -80ºC until further processing. Cells were lysed by bead beating with 3 sterile ceramic beads (Merck, Darmstadt, Germany) for 10 minutes, followed by total RNA isolation using the RNeasy Mini kit (Qiagen, Germantown, MD, USA) according to the manufacturer’s instructions. Samples were treated with two rounds of DNase to remove contaminating DNA using Turbo-DNase (Ambion). Messenger RNA (mRNA) libraries preparation and sequencing were performed by Novogene (Shanghai, PRC) using Novaseq 2x150 (Illumina, San Diego, USA).

***Diatom transcriptome analyses.*** Illumina reads were checked for quality using FastQC version v0.11.9 (https://www.bioinformatics.babraham.ac.uk/projects/fastqc/) and multiQC v1.11^14^ for all datasets. Adapter and low-quality reads were removed using fastp v0.23.2^15^. High-quality (Q score >30) reads were mapped to the *Asterionellopsis glacialis* A3 reference genome (GCA_014885115.2)^3^ using HISAT2 v2.1.0^16^ with default parameters. Alignment files in SAM format were converted into BAM format and then sorted by genomic coordinates using SAMtools v1.21^17^. Mapping statistics were generated using *flagstat* within samtools v1.21 to assess the number and percentage of mapped reads and properly paired alignments for each sample. Sorted BAM files were used for gene-level expression quantification using featureCounts from the Subread package v2.0.3^18,19^. Paired-end reads were counted as fragments using the “-p” and “--countReadPairs” options, and only fragments mapping to exons were considered. Gene-level counts were summarized using the gene name attribute in the annotation (GFF) file. The Trinity toolkit v2.15.1^20^ running inside a Singularity container was then employed to ensure that the biological replicates were well correlated, using the function “compare_replicates*”* for each condition. The obtained gene-level counts were used for differential expression analysis using the Trinity script “run_DE_analysis.pl*”* with the DESeq2 method^21^. Genes with a false discovery rate (FDR) threshold of 0.05 and |log₂ fold change| ≥ 1 were considered significantly differentially expressed. The diatom genome annotation (<https://zenodo.org/records/18194605>) was obtained using the automated Genome Sequence Annotation Server GenSAS^22^.

**Bacterial incubation experiments.** *P. pseudonitzschiae* grown in Zobell marine broth were monitored until an OD_600_ of 0.5 was reached. Cultures were then centrifuged at 4000 rpm for 10 min followed by washing twice with sterile 10% marine broth. To test the transcriptional response of *P. pseudonitzschiae* to 4-pyridoxolactone, pyridoxamine or pyridoxal or a mixture of all three metabolites (Merck, Darmstadt, Germany), washed cells were resuspended at a starting density of OD_600_ of 0.2 in 10% marine broth. Cultures were supplemented with a final concentration of 50 μM filter-sterilized solution of each metabolite individually or 16.6 μM of all three metabolites when combined. Controls were supplied with an equal volume of solvent necessary to dissolve the metabolite of interest. All experiments were conducted in triplicates.

***Bacterial RNA isolation and sequencing.*** For *P. pseudonitzschiae* RNA, cells were filtered through 0.2 µm polycarbonate filters (25 mm, Whatman, NJ, USA). All filters were flash frozen in liquid nitrogen and later stored in -80ºC until further processing. From the 0.2-µm filters, cells were lysed by bead beating with sterile ceramic beads (Merck, Darmstadt, Germany) for 10 minutes followed by total RNA isolation using the RNeasy Mini kit (Qiagen, German town, MD, USA) according to the manufacturer’s instructions. Samples were treated with two rounds of DNase to remove contaminating DNA using Turbo-DNase (Ambion). Ribosomal RNA (rRNA) were removed using the MicrobExpress Bacterial mRNA enrichment kit (Ambion) following the manufacturer’s instructions. RNA libraries preparation and sequencing were performed by Novogene (Shanghai, PRC) using Novaseq 2x150 (Illumina, San Diego, CA, USA).

***Bacterial transcriptome analyses.*** Reads were checked for quality as above. High-quality reads were mapped to the *P. pseudonitzschiae* F5 reference genome (GCF_014805345.1)^23^ using bowtie2 v2.3.3^24^ with default parameters. Alignment files were processed as described above to obtain sorted BAM files. These were used for gene-level expression quantification using featureCounts using the following parameters: "-t CDS” and “-g locus_tag" to get gene-level expression counts. Raw counts were used for differential expression analysis within Trinity and DESeq2 as described above. The same significance thresholds (FDR and fold change) were implemented.

***Functional enrichment analyses.*** KEGG pathway enrichment was performed using the *R* package clusterProfiler (v4.14.6)^25^. Custom KEGG Orthology (KO) annotations were used for both *A. glacialis* and *P. pseudonitzschiae*. Differentially upregulated genes at 60 and 180 hours were mapped to KO identifiers and tested for pathway enrichment against a background comprising all annotated KOs per species. Enrichment was assessed with the “enrichKEGG function” and organism set to “ko” for non-model species, and significance was determined using Benjamini–Hochberg–adjusted p-values (FDR < 0.05).

**Determination of Urea production.** *P. pseudonitzschiae* cells prepared as above were incubated with 4-pyridoxolactone at a final concentration of 50 μM for 24 hours after which cells were centrifuged at 4000 rpm for 10 mins. Urea production was determined using the Abcam Urea Assay Kit ab83362 (Cambridge, UK) according to manufacturer’s recommendations. For absorbance measurements at 570 nm, an Epoch plate reader (BioTek, Winooski, USA) was used.

***Determination of protein function.*** To identify putative matches for PLDH and PDLA, the auto-annotation of the *A. glacialis* A3 and *P. pseudonitzschiae* F5 genomes were mined for enzymes belonging to the short-chain dehydrogenase/reductase activity (SDR) family and metallo beta-lactamase (MBL) family, respectively^26^. The protein structures for all matches in those families were modeled using Alphafold2.0^27^ and aligned with known protein crystal structures of *Mesorhizoium loti* (3RWB)^26^ and *M. japonicum* (4KEQ)^28^ using RCSB pairwise align (<https://www.rcsb.org/alignment>)^29^. Candidates were chosen according to RMSD, TM-Score, query vs subject length, and residue identity (Supplementary Information Table 20-21). Further active sites (substrate binding sites) were investigated using Pymol 3.1.3 (<http://www.pymol.org/pymol>) to compare potential active site mutations. Diatom and bacterial protein candidates were excluded according to mismatches in amino acids residues within the binding sites. For the diatom, A3_.00g105710.m01 and A3_.00g133870.m01 fit the criteria we used and for the bacterium, only IHQ53_RS05450 did so. These protein targets were used for functional testing.

Proteins were expressed using the protein expression service from Genscript (Piscataway, USA). In short, for recombinant protein expression genes were codon optimized, modified with His-Tag and expressed in a pET30a vector in *E. coli* strain BL21 Star™ (DE3) for 16h in 4°C or 4h at 37°C. Expressed proteins were purified over a Ni-column and supplied in 50 mM Tris-HCl, 150 mM NaCl, 10% Glycerol, pH 8.0. Expressed and purified proteins were recovered from the soluble fraction apart from the bacterial protein, which was additionally recovered and refolded from inclusion bodies. For refolding the supplied expressed proteins were denatured in 6M urea and transferred into dialysis cassettes (Slide-A-Lyzer, ThermoFischer Scientific, Waltham, USA)^30^. Refolding was performed by step wise decrease of urea concentration through dialysis after initial incubation at 6 M for 2h, 4 M (for 4h, 2 M (for 4h) to 0 M (overnight) in 50 mM Tris-HCl, 50 mM and 150 mM NaCl buffer at room temperature. Additional Zn^2+^ was added at a final concentration of 10 μM for the pyridoxolactonase candidate. After refolding the protein concentrations were determined using the Qubit fluorometric protein assay kit (Invitrogen, Waltham, USA) according to manufacturer’s recommendations.

***Enzyme activities.*** Activities of the three expressed proteins were tested using UV absorption to detect the concentration of 4-pyridoxolactone. For pyridoxal dehydrogenase, enzyme activity was measured at 25°C in 1 mL of a reaction mixture consisting of 50 mM Tris-HCl, 150 mM NaCl (pH 7.2), 1 mM pyridoxal, 1 mM NAD^+^, and 1 µM enzyme. The molecular extinction coefficient of 4-pyridoxolactone (7500 M^−1^ cm^−1^ ) was used to calculate the product concentrations at 340 nm after correcting for NADH^+^ absorption (6600 M^−1^ cm^−1^)^28,31^. For 4-Pyridoxolactonase, enzyme activity was assayed by following the decrease in absorbance at 340 nm, indicating the hydrolysis of 4-pyridoxolactone to 4-pyridoxate. The 1 mL assay mixture consisted of PBS buffer (pH 7.2), 0.25 mM ZnCl_2_, 0.5 mM 4-pyridoxolactone and 4-Pyridoxolactonase at a final concentration of 0.5 mM. Absorbance was determined on an Epoch plate reader (BioTek, Winooski, USA) using 100 μL of the reaction mixture in flat bottom 96 well UV-star microplates (Greiner Bio-One, Kremsmuenster, AUT).

***Global distribution of PLDH and PDLA in the Oceans and in public genomes.*** The protein sequences of PLDH from *A. glacialis*, *M. loti,* and *M. japonicum*, and PDLA from *P. pseudonitzschiae* and *M. japonicum* were used to obtain hidden Markov model (HMM) profiles with the hmmbuild function within HMMER v3.4. The generated HMM profiles were used as queries against metagenomes and metatranscriptomes datasets of the Tara Oceans Gene Atlas (http://tara-oceans.mio.osupytheas.fr/ocean-gene-atlas/) web server^32^ with an e-value threshold of 1e-20. We inferred potential signaling when both genes or transcripts were present/expressed simultaneously at the same depth and station, irrespective of the size fraction. We also only included samples from the surface (SRF) and deep chlorophyll maximum (DCM) in Tara and did not mine other depths since phytoplankton are restricted to the sunlit ocean. Geographic distributions and taxonomy assignments in Tara metagenomes were visualized as pie charts across a world map at the surface and the deep chlorophyll maximum using R packages ggplot2 v4.0.1, sf v1.0-23, rnaturalearth v1.1.0, and scatterpie v0.2.6. The transcript relative abundance in Tara metatranscriptomes was plotted. The same HMM profiles were then queried against the RefSeq database (https://www.ncbi.nlm.nih.gov/refseq/) using hmmscan with an e-value threshold of 1e-20 for PDLA and 1e-50 for PLDH. Duplicate hits and hits with <200 amino acids were discarded. The remaining sequences were clustered using USEARCH^33^ with 90% identity. The resulting sequences, in addition to the ones used to build the HMM profiles, were aligned on MUSCLE v3.8.31 for PDLA and MAFT v7.526 for PLDH. The alignments were then trimmed using trimAl v1.5.1^34^ on "gappyout" mode. Phylogeny was inferred using FastTree v2.2^35^. The resulting newik trees were visualized on the Interactive Tree of Life (iTOL) tool v6^36^.

**References**

1. Ryther, J. H. & Guillard, R. Studies of marine planktonic diatoms: II. Use of Cyclotella nana Hustedt for assays of vitamin B12 in sea water. *Canadian Journal of Microbiology* **8**, 437–445 (1962).

2. Brand, L. E., Guillard, R. R. L. & Murphy, L. S. A method for the rapid and precise determination of acclimated phytoplankton reproduction rates. *Journal of Plankton Research* **3**, 193–201 (1981).

3. Shibl, A. A. *et al.* Diatom modulation of select bacteria through use of two unique secondary metabolites. *Proc National Acad Sci* **117**, 27445–27455 (2020).

4. ZoBell, C. E. Studies on marine bacteria. I. The cultural requirements of heterotrophic aerobes. *J. mar. Res* **4**, 42–75 (1941).

5. Polerecky, L. *et al.* Look@NanoSIMS – a tool for the analysis of nanoSIMS data in environmental microbiology. *Environ. Microbiol.* **14**, 1009–1023 (2012).

6. Kitzinger, K. *et al.* Fluorescence In-Situ Hybridization (FISH) for Microbial Cells, Methods and Concepts. *Methods Mol. Biol.* **2246**, 207–224 (2021).

7. Foster, R. A. *et al.* The rate and fate of N2 and C fixation by marine diatom-diazotroph symbioses. *Isme J* 1–11 (2021) doi:10.1038/s41396-021-01086-7.

8. Ochsenkühn, M. A., Schmitt-Kopplin, P., Harir, M. & Amin, S. A. Coral metabolite gradients affect microbial community structures and act as a disease cue. *Commun. Biology* **1**, 184 (2018).

9. Marshall, A. G., Hendrickson, C. L. & Jackson, G. S. Fourier transform ion cyclotron resonance mass spectrometry: A primer. *Mass Spectrom. Rev.* **17**, 1–35 (1998).

10. Kujawinski, E. B. & Behn, M. D. Automated Analysis of Electrospray Ionization Fourier Transform Ion Cyclotron Resonance Mass Spectra of Natural Organic Matter. *Anal. Chem.* **78**, 4363–4373 (2006).

11. Rahiman, N., Ochsenkühn, M. A., Amin, S. A. & Gunsalus, K. C. MIMI: Molecular Isotope Mass Identifier for stable isotope-labeled Fourier transform ultra-high mass resolution data analysis. *BMC Bioinform.* In press (2026) doi:10.1186/s12859-025-06348-1.

12. Djoumbou-Feunang, Y. *et al.* CFM-ID 3.0: Significantly Improved ESI-MS/MS Prediction and Compound Identification. *Metabolites* **9**, 72 (2019).

13. Weininger, D. SMILES, a chemical language and information system. 1. Introduction to methodology and encoding rules. *J. Chem. Inf. Comput. Sci.* **28**, 31–36 (1988).

14. Ewels, P., Magnusson, M., Lundin, S. & Käller, M. MultiQC: summarize analysis results for multiple tools and samples in a single report. *Bioinformatics* **32**, 3047–3048 (2016).

15. Chen, S., Zhou, Y., Chen, Y. & Gu, J. fastp: an ultra-fast all-in-one FASTQ preprocessor. *Bioinformatics* **34**, i884–i890 (2018).

16. Kim, D., Paggi, J. M., Park, C., Bennett, C. & Salzberg, S. L. Graph-based genome alignment and genotyping with HISAT2 and HISAT-genotype. *Nat. Biotechnol.* **37**, 907–915 (2019).

17. Li, H. *et al.* The sequence alignment/map format and SAMtools. *Bioinformatics* **25**, 2078–2079 (2009).

18. Liao, Y., Smyth, G. K. & Shi, W. featureCounts: an efficient general purpose program for assigning sequence reads to genomic features. *Bioinformatics* **30**, 923–930 (2014).

19. Liao, Y., Smyth, G. K. & Shi, W. The Subread aligner: fast, accurate and scalable read mapping by seed-and-vote. *Nucleic Acids Res.* **41**, e108–e108 (2013).

20. Haas, B. J. *et al.* De novo transcript sequence reconstruction from RNA-seq using the Trinity platform for reference generation and analysis. *Nat. Protoc.* **8**, 1494–1512 (2013).

21. Love, M. I., Huber, W. & Anders, S. Moderated estimation of fold change and dispersion for RNA-seq data with DESeq2. *Genome Biol.* **15**, 550 (2014).

22. Humann, J. L., Lee, T., Ficklin, S. & Main, D. Gene Prediction, Methods and Protocols. *Methods Mol. Biol.* **1962**, 29–51 (2019).

23. Fei, C. *et al.* Quorum sensing regulates ‘swim‐or‐stick’ lifestyle in the phycosphere. *Environ. Microbiol.* **22**, 4761–4778 (2020).

24. Langmead, B. & Salzberg, S. L. Fast gapped-read alignment with Bowtie 2. *Nat. Methods* **9**, 357–359 (2012).

25. Yu, G., Wang, L.-G., Han, Y. & He, Q.-Y. clusterProfiler: an R Package for Comparing Biological Themes Among Gene Clusters. *OMICS: A J. Integr. Biol.* **16**, 284–287 (2012).

26. Maatouk, M. *et al.* Metallo-Beta-Lactamase-like Encoding Genes in Candidate Phyla Radiation: Widespread and Highly Divergent Proteins with Potential Multifunctionality. *Microorganisms* **11**, 1933 (2023).

27. Jumper, J. *et al.* Highly accurate protein structure prediction with AlphaFold. *Nature* **596**, 583–589 (2021).

28. Yokochi, N., Nishimura, S., Yoshikane, Y., Ohnishi, K. & Yagi, T. Identification of a new tetrameric pyridoxal 4-dehydrogenase as the second enzyme in the degradation pathway for pyridoxine in a nitrogen-fixing symbiotic bacterium, Mesorhizobium loti. *Arch. Biochem. Biophys.* **452**, 1–8 (2006).

29. Bittrich, S., Segura, J., Duarte, J. M., Burley, S. K. & Rose, Y. RCSB protein Data Bank: exploring protein 3D similarities via comprehensive structural alignments. *Bioinformatics* **40**, btae370 (2024).

30. Yamaguchi, H. & Miyazaki, M. Refolding Techniques for Recovering Biologically Active Recombinant Proteins from Inclusion Bodies. *Biomolecules* **4**, 235–252 (2014).

31. McCormick, D. B. Vitamin B6. in *Methods in Enzymology* vol. 122 93–103 (Academic Press, Orlando, FL, 1986).

32. Vernette, C. *et al.* The Ocean Gene Atlas v2.0: online exploration of the biogeography and phylogeny of plankton genes. *Nucleic Acids Res.* **50**, W516–W526 (2022).

33. Zhou, Y., Liu, Y. & Li, X. USEARCH 12: Open‐source software for sequencing analysis in bioinformatics and microbiome. *iMeta* **3**, e236 (2024).

34. Capella-Gutiérrez, S., Silla-Martínez, J. M. & Gabaldón, T. trimAl: a tool for automated alignment trimming in large-scale phylogenetic analyses. *Bioinformatics* **25**, 1972–1973 (2009).

35. Price, M. N., Dehal, P. S. & Arkin, A. P. FastTree 2 – Approximately Maximum-Likelihood Trees for Large Alignments. *PLoS ONE* **5**, e9490 (2010).

36. Letunic, I. & Bork, P. Interactive Tree of Life (iTOL) v6: recent updates to the phylogenetic tree display and annotation tool. *Nucleic Acids Res.* **52**, W78–W82 (2024).
